# Dynamic evolution of satellite DNAs drastically differentiates the genomes of *Tribolium* sibling species

**DOI:** 10.1101/2024.12.02.626372

**Authors:** Damira Veseljak, Evelin Despot-Slade, Marin Volarić, Lucija Horvat, Tanja Vojvoda Zeljko, Nevenka Meštrović, Brankica Mravinac

## Abstract

Tandemly repeated satellite DNAs (satDNAs) are among the most abundant and fastest-evolving eukaryotic sequences, but the way they model genomes is still elusive. Here, we investigated the evolutionary dynamics of satDNAs in the extremely satDNA-rich genomes of two closely related *Tribolium* insects that produce sterile hybrids. In *Tribolium freemani*, we identified 135 satDNAs, accounting for 38.7% of the genome. Comparative analysis with the *Tribolium castaneum* satellitome revealed that the drastic difference happened in their centromeric regions, which share orthologous organization hallmarked by totally different major satDNAs but related minor satDNAs. The *T. freemani* male sex chromosome, which lacks the major satDNA but contains a minor-like satDNA, further heightened the question of which satDNA is centromere-competent. By analyzing the long-range organization of the centromeric regions, we revealed that both the major and minor satDNA arrays exhibit a strong tendency toward macro-dyad symmetry, suggesting that the secondary structures in the centromeres may be more important than the primary sequence itself. We found evidence that the centromeric satDNAs of *T. freemani* occur in extrachromosomal circular DNAs, which may contribute to their expansion and homogenization between non-homologous chromosomes. We also identified numerous low-copy-number satDNAs that are orthologous between the siblings, some of which are associated with transposable elements, highlighting transposition as a mechanism of their spreading. The dynamic evolution of satDNAs has clearly influenced the differentiation of *Tribolium* genomes, but the question remains whether the differences in their satDNA profiles are a cause or consequence of speciation.

## INTRODUCTION

Satellite DNAs (satDNAs) are highly repetitive, tandemly organized DNA sequences whose contiguous arrays make up extensive regions in many eukaryotic genomes. The repetitive organization has hampered their study in the past and made them the most challenging regions to assemble in sequenced genomes. They were initially considered “junk DNA” due to their abundance and unclear function, but the accumulation of studies on numerous organisms began to change their unfavorable reputation over time. Since they frequently form heterochromatin blocks in the (peri)centromeric regions of many eukaryotes, their most recognized role is attributed to the organization of functional centromeres (Hartley and O’Neill 2019; Talbert and Henikoff 2022), which is consequently associated with proper chromosome segregation and genome stability (Flynn and Yamashita 2024). Aberrant transcription of some pericentromeric satDNAs has been associated with a tumor-promoting function (Bersani et al. 2015; Iwata et al. 2024). The evolutionary significance of satDNAs is supported by evidence that these sequences may be involved in speciation (Ferree and Barbash 2009; Jagannathan and Yamashita 2021). However, despite various ascribed roles, no satDNA function has yet been found to be ubiquitous in all organisms studied.

SatDNAs are among the most rapidly evolving eukaryotic sequences, and can differ significantly in nucleotide sequence and copy number of their repeat units even between related species. The main theories explaining the evolution of satDNAs were proposed at a time when, due to methodological limitations, only the most abundant satellites were discovered in genomes. According to concerted evolution, the gradual accumulation of mutations between repeat units of a given satellite leads to greater interspecific repeat variability compared to intraspecific variability (Elder and Turner 1995). In some cases, large differences between various satDNAs that cannot be explained by the process of gradual degradation of the nucleotide sequence are explained by the satDNA library theory. The library theory implies the existence of a satellite collection of different satDNAs in an ancestral genome, of which certain satDNAs significantly change the number of copies by amplification or reduction, leading to different satellite profiles in descendant species (Fry and Salser 1977; Meštrović et al. 1998). Modern sequencing technologies and accompanying bioinformatics tools have improved the detectability of satDNAs, so that often up to over a hundred satDNAs can be detected in the genomes of different organisms (Boštjančić et al. 2021; João Da Silva et al. 2023). The availability of an increasing number of high-throughput satellitome analyses contributes to the understanding of satDNA evolution and encourages the revision of previous evolutionary concepts (Belyayev et al. 2020). For these purposes, comparative analyses of related species are very informative (Camacho et al. 2022), whereby organisms with a high proportion of satDNAs represent an additional challenge, but also a beneficial source of information.

The flour beetle *Tribolium freemani* belongs to the genus that includes some of the most important pests of stored agricultural products (Sokoloff 1972). In addition to their economic importance, *Tribolium* species are also an excellent platform for the study of satellite DNAs, which occupy up to 60% of their genomes (Mravinac and Plohl 2010). The representative of the genus *Tribolium*, but also of the entire order Coleoptera, is *Tribolium castaneum*. It was the first beetle whose genome was sequenced in 2008 (Richards et al. 2008), and the assembly has been refined and improved in the meantime (Herndon et al. 2020), with the latest assembly version TcasONT being significantly enhanced in the repetitive part of the genome (Volarić et al. 2024). The satDNA studies have revealed that the *T. castaneum* genome contains at least 57 different satellites (Ugarković, Podnar, et al. 1996; Feliciello et al. 2015; Pavlek et al. 2015; Gržan et al. 2023), of which the major satellite TCAST comprises 17% of the genome. The closest congener to *T. castaneum* is *T. freemani*. The two species are so closely related that they can hybridize but produce sterile offspring (Nakakita et al. 1981). We recently sequenced and assembled the *T. freemani* genome and found that the two siblings are very similar in their coding sequence (Volarić et al. 2022). Regarding satDNAs, it was discovered three decades ago that 31% of the *T. freemani* genome is made of one satDNA (Juan et al. 1993), whose nucleotide sequence shows absolutely no similarity to the major satellite of *T. castaneum,* suggesting an independent origin. However, in both species the major satellites, although remarkably different in sequence, occupy the (peri)centromeric regions (Juan et al. 1993; Ugarković, Podnar, et al. 1996; Gržan et al. 2020).

Understanding the evolutionary dynamics by which satDNAs mold genomes is necessary for the perception of genome versatility, and it may also be an essential step in unraveling the role of these functionally controversial sequences. Considering the extremely variable nature of satDNAs and inspired by the fact that *T. freemani* and *T. castaneum* produce infertile hybrid progeny, in this work we address the question of how compatible the two species are in their satellite profiles. In an initial satellitome analysis, we discovered 135 satDNAs in *T. freemani* by combining an assembly-free detection and an assembly-based inspection. By revising the major satellite and analyzing its ultra-long arrangement, we deciphered the organization of the centromeric regions and revealed which satDNAs are involved in the structure of these architecturally and functionally crucial chromosomal parts. We also discovered the origin of diverse major satDNAs of *T. freemani* and *T. castaneum*. Finally, by exploring the orthologous satDNAs of the two species, we draw conclusions about the evolutionary dynamics and the main molecular mechanisms that most apparently differentiated the siblings’ satellitomes and their genomes in general.

## RESULTS

### 1. Identification of *T. freemani* satDNA repeats

To decipher the *T. freemani* satellitome by using an assembly-independent approach, we re-sequenced its genome with Illumina sequencing. Randomly subsampled sets of Illumina reads corresponding to different genome coverages were subjected to graph-based clustering using the TAREAN program (**Suppl. Table S1**). Consensus sequences of the clusters annotated as potential satDNA were mapped to the *T. freemani* reference genome assembly Tfree1.0 (Volarić et al. 2022), applying the criterion of ≥70% sequence similarity. The candidate sequences that were mapped in ≥5 consecutive copies were declared satDNAs. In this way, 135 satDNAs were identified, one already known from previous studies (Juan et al. 1993) and 134 new ones. The consensus sequences of the 135 satDNAs, named TfSat01-135, are listed in the **Suppl. Table S2**.

Of the 135 discovered satDNAs, 124 were found to be unrelated, while 11 showed partial similarity (56.7-73.0%) in nucleotide sequence, suggesting a related origin. Based on the similarities, we classified the related satDNAs into the four superfamilies, SF1-SF4 (**Suppl. Fig. S1, Suppl. Table S3**). The consensus sequences of the new satDNAs were also searched against the NCBI GenBank nucleotide database. The only similarity found for some satDNAs was with previously described satellites of the sibling species *T. castaneum* (**Suppl. Table S3**). When compared with repetitive sequences from the Repbase collection, the 75 *T. freemani* satDNAs showed only segmental similarities with different mobile elements, mainly DNA transposons (**Suppl. Table S3**).

Regarding repeat unit lengths, the consensus sequences of the 135 satDNAs range from 63 bp to 1106 bp (**Suppl. Table S3**), with an evident preference for lengths between 140-180 bp (**Fig. 1A**). For five satDNAs, we found that their monomers are higher-order repeats (HOR) based on two or three 90-188 bp long subunits, sharing 63.0-82.2% pairwise similarity (**Suppl. Fig. S2**). Another feature that largely characterizes the *T. freemani* satellitome is the biased DNA base composition, with 127 satDNAs having an A+T content >60% (**Fig. 1A**, **Suppl. Table S3**).

**Figure 1.**
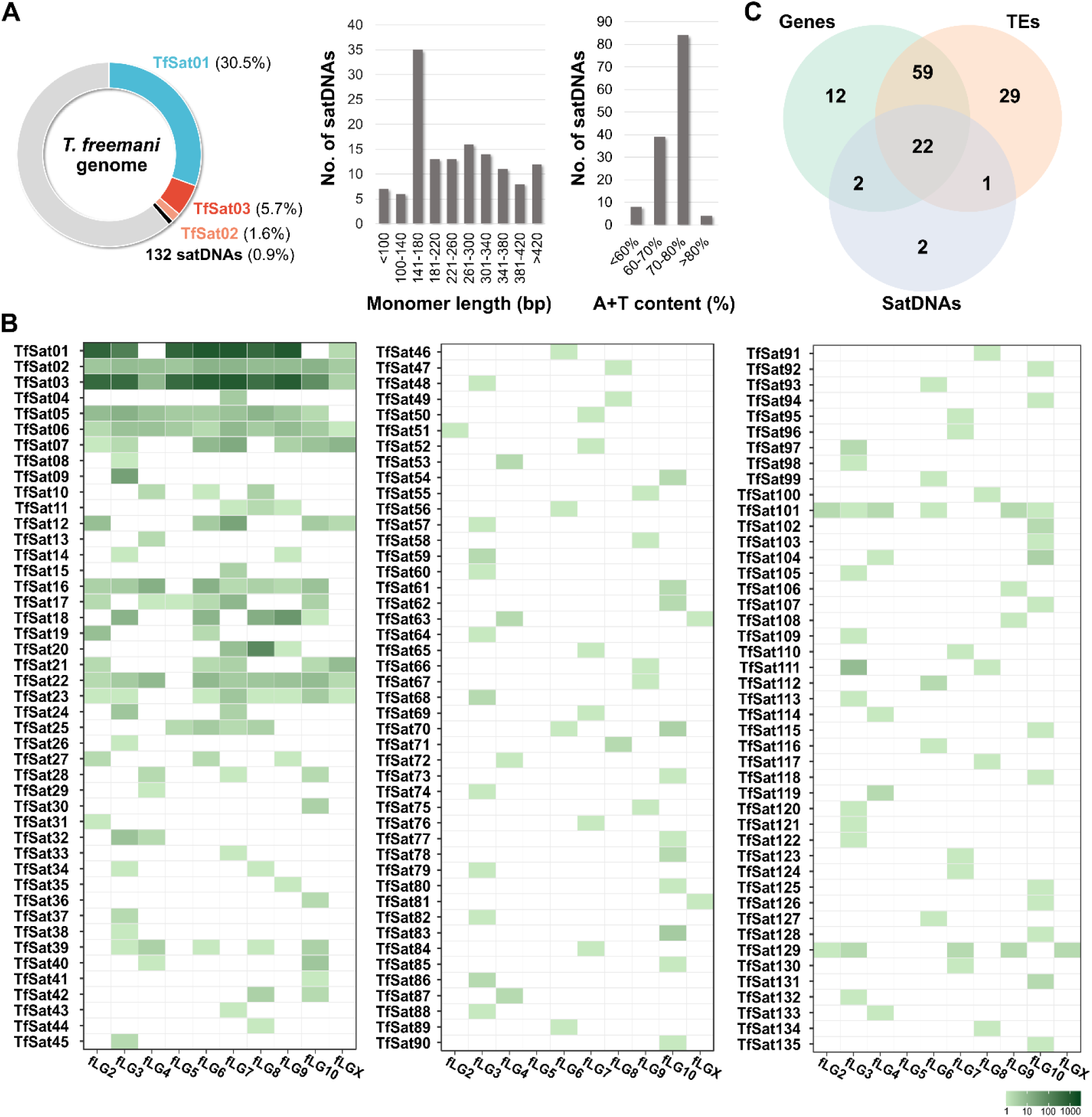
The overview of the *T. freemani* satellitome comprising 135 satDNAs. **A)** The genomic proportions of the three most abundant satDNAs and 132 low-copy-number satDNAs, and their distributions regarding the monomer length and A+T content of the consensus sequences. The genomic proportions are based on the estimates according to the TAREAN analyses, while the distributions of monomer lengths and A+T composition were derived from the data for the individual satDNAs listed in Supplementary Table S3. **B)** The presence of satDNA arrays with ≥5 consecutive copies mapped on the ten *T. freemani* chromosomes, fLG2-fLGX, in the genome assembly Tfree1.0. The number of annotated copies on chromosomes is color-coded according to the color scale. **C)** Venn diagram showing the number of the *T. freemani* satDNAs that have annotated genes, transposable elements (TEs) and/or satDNAs in the 10-kb regions flanking their arrays. The diagram is based on the data for each satDNA shown in Supplementary Table S6.

The satellitome with the 135 satDNAs comprises 38.7% of the *T. freemani* genome (**Fig. 1A**). However, the genome is dominated by a single, extremely abundant satDNA, TfSat01, which makes up almost one third of the genome sequence. In addition, the two satDNAs, TfSat02 and TfSat03, are moderately represented (1.6% and 5.7%, respectively), while the remaining 132 satDNAs together account for only 0.9% of the genome and can be considered low-copy-number satellites (**Fig. 1A**, **Suppl. Table S3**). Regarding the distribution, the repeats of the three most abundant satDNAs were detected on almost all chromosomes, whereas the low-copy-number satDNAs showed different distribution patterns (**Fig. 1B, Suppl. Fig. S3**). The majority of them (101 satDNAs) were assigned only to one chromosome in the Tfree1.0 assembly, and a smaller number of them were mapped to multiple chromosomes (**Fig. 1B**). To verify the chromosomal distribution of the low-copy-number satellites obtained *in silico*, we experimentally analyzed six of them with a genome proportion between 0.01% and 0.05%. We found that satDNAs with a genome proportion higher than 0.02% had a broader chromosomal distribution (**Suppl. Fig. S3A-C**), while satDNAs with a lower genome proportion (<0.02%) revealed a distinct signal only on one chromosome pair (**Suppl. Fig. S3D-E**), which is generally consistent with the *in silico* analysis. Further, regarding the repetitiveness of monomeric units, repeats of 71 satDNAs were found exclusively in arrays containing ≥5 consecutive monomers, while for 64 satDNAs, in addition to their longer arrays, we also detected the shorter stretches with less than 5 consecutive copies (**Suppl. Table S4**).

To gain insight into the genomic environment of the identified satDNAs, we analyzed the 10 kb regions upstream and downstream of the satDNA arrays for the presence of other satDNAs as well as transposable elements (TEs) and genes annotated in the Tfree1.0 assembly. It turned out that 111 satDNAs have TEs, 95 have genes, and only 27 of them have other satellite sequences in their 10-kb flanking regions **(Fig. 1C, Suppl. Table S5-S6, Suppl. Fig. S4**). The concurrent presence of TEs and genes was detected in the surrounding regions of as many as 59 satDNAs, while only 22 satDNAs harbor TEs, genes, and satDNAs in their 10-kb proximity. Even when present, the satDNAs generally did not stand out as significantly closer neighbors. Namely, regarding the distance of a nearest annotated element to a satDNA array, similar average distances of 4-5 kb were found for all three addressed sequence types (genes, satDNAs, TEs) (**Suppl. Table S7**). Therefore, we conclude that the *T. freemani* satellitome is not exclusively associated with gene-poor regions and that most low-copy-number satellites are not located in regions dominated by tandemly repeated sequences.

The obtained general overview of the satellitome served as a starting point for detailed analyses of the most prominent satDNAs in the *T. freemani* genome and the investigation of the evolutionary dynamics and molecular mechanisms that shaped the satellitome.

### 2. TfSat01, the main constituent of the *T. freemani* satellitome and centromere

#### Revision of the major satDNA TfSat01

First, we reexamined the most plentiful satDNA, named TfSat01 in this study. The 166 bp long TfSat01 sequence corresponds to the previously described major satDNA, which was experimentally determined to comprise 31% of the *T. freemani* genome (Juan et al. 1993), also supported by our TAREAN analyses (**Suppl. Table S3**). However, our TfSat01 consensus, based on 228,060 monomers annotated in the Tfree1.0 genome assembly, showed 5.4% nucleotide difference compared to the 30-year-old GenBank entry X58539, the consensus derived from five cloned monomers (Juan et al. 1993) (**Suppl. Fig. S5A**). We hold that the TfSat01 consensus better represents the *T. freemani* major satDNA (argued in detail in the Material and Methods section), so we used it in all downstream analyses.

The major satDNA was previously reported to be located in centromeric chromosomal areas of *T. freemani* (Juan et al. 1993). *T. freemani* has a typical coleopteran karyotype, 2n=20, with nine pairs of autosomes and one pair of sex chromosomes (Shimeld 1989). The sex chromosome pair in females consists of two X chromosomes, while in males there is an Xy_p_ parachute-like association of the X and a tiny y_p_ chromosome. To verify TfSat01 presence on all chromosomes, we performed fluorescence *in situ* hybridization (FISH) on female and male metaphase chromosome spreads (**Fig. 2A**). We indeed found TfSat01 present on all 20 chromosomes in females. In males, however, we detected TfSat01 signals on 18 autosomes and the X chromosome, but surprisingly not on the male-specific y_p_ chromosome, revealing that the sex chromosome y_p_ is deprived of the major satDNA TfSat01 (**Fig. 2A**).

**Figure 2.**
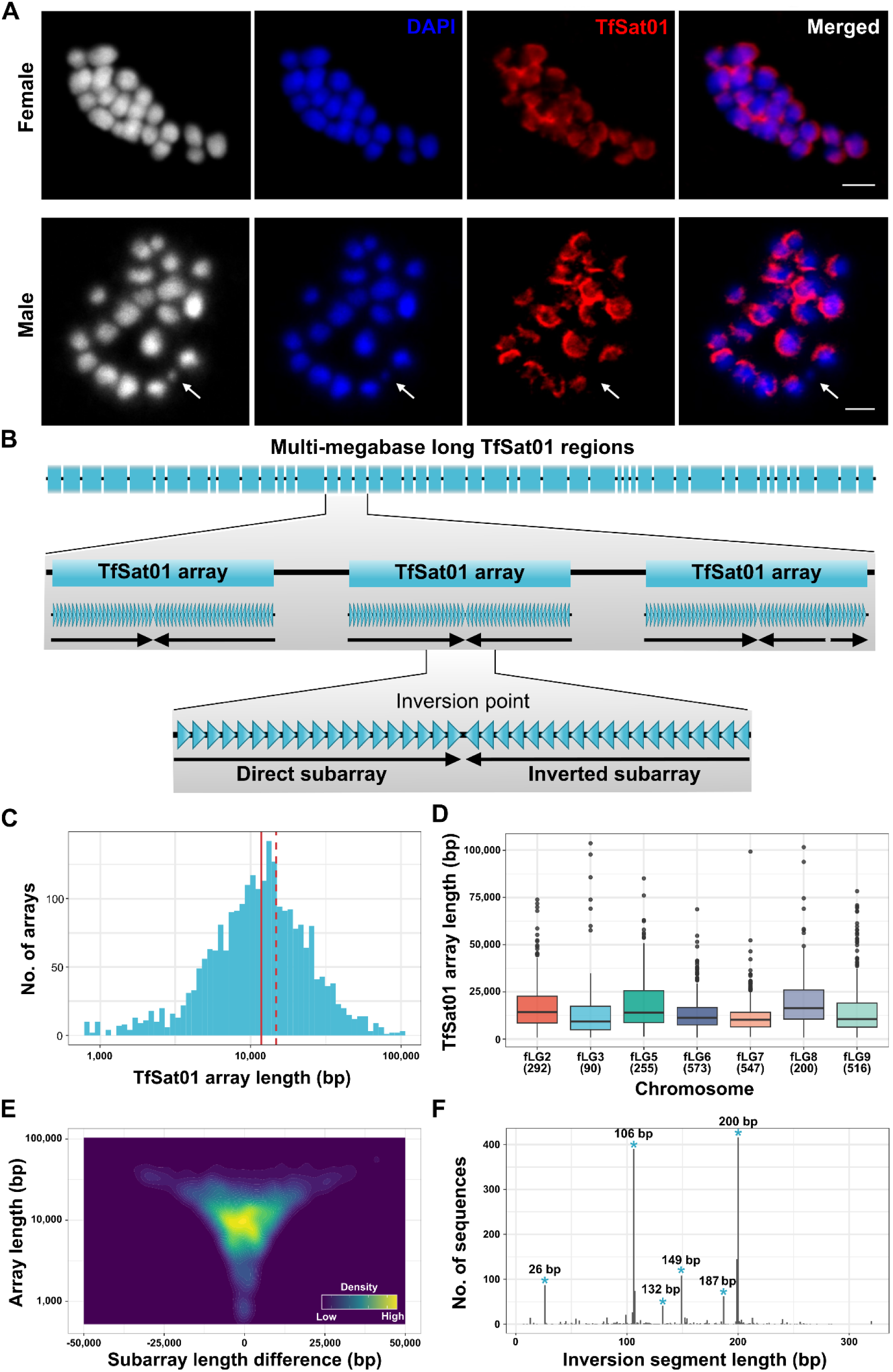
The organization of the major satDNA TfSat01 in the *T. freemani* genome. **A)** Localization of TfSat01 satDNA on the *T. freemani* female (2n=18+XX) and male (2n=18+Xy_p_) metaphase chromosomes determined by fluorescence *in situ* hybridization. The first panels show the chromosomes in a black and white version to better visualize the contours of the chromosomes, especially the male minute chromosome y_p_. The chromosomes are stained in DAPI (blue fluorescence), while TfSat01 signals are shown in red fluorescence. A white arrow points to the y_p_ chromosome lacking the TfSat01 signal. The bar represents 3 µm. **B)** A schematic illustrating the long-range organization of the multi-megabase long regions consisting of TfSat01 arrays. Within the TfSat01 array, the TfSat01 monomers (blue triangles) are repeatedly organized into subarrays that differ from each other by the orientation of the monomers that form them. Thus, the term “TfSat01 array” refers to a continuous array of TfSat01 monomers, regardless of the number of subarrays it contains. **C)** Distribution of the lengths of the TfSat01 arrays. The red solid line represents the median (11.8 kb), while the mean (14.8 kb) is indicated by the dashed line. **D)** The box plot analysis of TfSat01 array length distribution at different *T. freemani* chromosomes. The black line within a box represents the median length, and the number of analyzed arrays is indicated in parentheses below the chromosome name. **E)** Density plot of the differences between the length of the direct and inverted subarrays within the 1793 TfSat01 arrays. The x-axis shows the difference in length between directly and inversely oriented subarrays within an array. The relative abundance of subarray length differences in the graph is indicated by a color gradient. Density plots for arrays on individual chromosomes are presented in Supplementary Figure S7. **F)** The length distribution of the inversion segments from the 1793 TfSat01 arrays. The blue dots in the graph indicate the six groups of the most frequent inversion segments, whose consensus sequences and detailed alignments are shown in Supplementary Figure S8 and Supplementary Data S1, respectively.

#### Long-range organization of the major satDNA TfSat01

Given that one third of the genome consists of TfSat01 repeats, we wanted to investigate their large-scale arrangement. The Tfree1.0 genome assembly lacks the assembled TfSat01 arrays on chromosomes fLG4, fLG10 and fLGX (Volarić et al. 2022), but multi-megabase TfSat01 arrays are successfully assembled on seven other chromosomes, which allowed us to explore their long-range organization.

Using the StainedGlass tool we visualized these multi-megabase regions, which span from 3.8 to 9.7 Mb (**Suppl. Table S8).** Sequence identity heatmaps revealed that these regions consist predominantly of arrays showing a high degree of identity (>93) **(Suppl. Fig. S6)**. However, at higher resolution, TfSat01 regions showed a more complex structure than a monotonous repetition of basic units (**Fig. 2B**). We determined that the median length of uninterrupted TfSat01 arrays is 11.8 kb (**Fig. 2C**). While the number of mapped TfSat01 arrays varies by up to 6.4-fold across different chromosomes, the median length of continuous arrays remains rather consistent, ranging from 9.3 to 16.3 kb (**Fig. 2D**). This suggests that TfSat01 array lengths are generally balanced throughout the genome.

Interestingly, the TfSat01 arrays have a specific substructure consisting of two or more subarrays that show inverted orientation of the TfSat01 repeats (**Fig 2B**). TfSat01 arrays with two inversely oriented subarrays are the most numerous (71.8% of all mapped arrays). Among the remaining arrays, those with an even number of inversely oriented subarrays predominate over those with an odd number (**Suppl. Table S9**). The favored even number of subarrays within most TfSat01 arrays may suggest a tendency toward dyad symmetry. An analysis of hypothetic hairpin structures in the 1793 arrays consisting of two subarrays showed that the hairpins would have some degree of asymmetry in their stems due to the varying number of TfSat01 monomers in the direct and inverted subarrays. The lack of perfect symmetry indicates potential flexibility in the organization of the subarrays. We found that the length difference between direct and inverted subarrays increases with array length (**Fig. 2E**), a correlation that holds across all chromosomes (**Suppl. Fig. S7**).

Next, we focused on the inversion segments where the subarrays change orientation. We discovered that the majority of these inversion segments fall into one of six groups defined by a specific length (26 bp, 106 bp, 132 bp, 149 bp, 187 bp, and 200 bp; **Fig. 2F**, **Suppl. Fig. S8A-B**). All six inversion segment types are based on an abruptly terminated TfSat01 monomer followed by a reverse-oriented TfSat01 monomer truncated at a different nucleotide position (**Suppl Fig. S8A-B**). Despite different truncation sites, all six inversion segment types have two characteristics in common: 1) there are no other extraneous intervening sequences between the reverse-oriented truncated TfSat01 copies, and 2) the reverse-oriented truncated copies do not exhibit increased nucleotide sequence variability (**Suppl. Fig. S8B**). Being reverse complemented, the truncated copies extend the stems of potential hairpins almost to the tip, ending in 9–27 bp long loops (**Suppl. Fig. S8C**). Importantly, these six inversion segment types are not chromosome-specific, as each of them was detected on three to seven chromosomes (**Suppl. Fig. S8D).** Moreover, the sequences of each inversion type are highly conserved, with an average similarity of 95.0%-99.9%. In fact, 100% identical copies of each inversion type were identified on non-homologous chromosomes (**Suppl. Data S1**). From this we conclude that the dyad symmetries and potential secondary structures formed by the TfSat01 arrays are conserved features of the *T. freemani* centromeric regions.

### 3. TfSat03 resides in the intercalary segments between the TfSat01 arrays

Our further analysis of TfSat01 long-range organization revealed that the intervals between TfSat01 arrays tend to be around 4 kb (**Fig. 3A-B**). We discovered that these intercalary segments mainly harbor repeats of TfSat03, the second most abundant satDNA, whose 340 bp long monomers make up 5.7% of the *T. freemani* genome (**Fig. 1A, Suppl. Table S3**). While TfSat03 can form long continuous arrays, such as a 41 kb region on chromosome fLG3, it predominantly resides in the intercalary segments between TfSat01 arrays. These intercalary segments are structured as “cassettes” (**Fig. 3A**) containing: 1) a variable number of highly conserved TfSat03 copies, typically between 6-8 (**Fig. 3C**), 2) degenerate TfSat03 repeats that are truncated and/or show <80% similarity to the consensus, and 3) symmetrically arranged 200-300 bp long outward-facing stretches abundant in (AAT)_n_ microsatellite (**Fig. 3A**). It is significant that these elements together form dyad symmetry within the intercalary segments (**Fig. 3A**).

**Figure 3.**
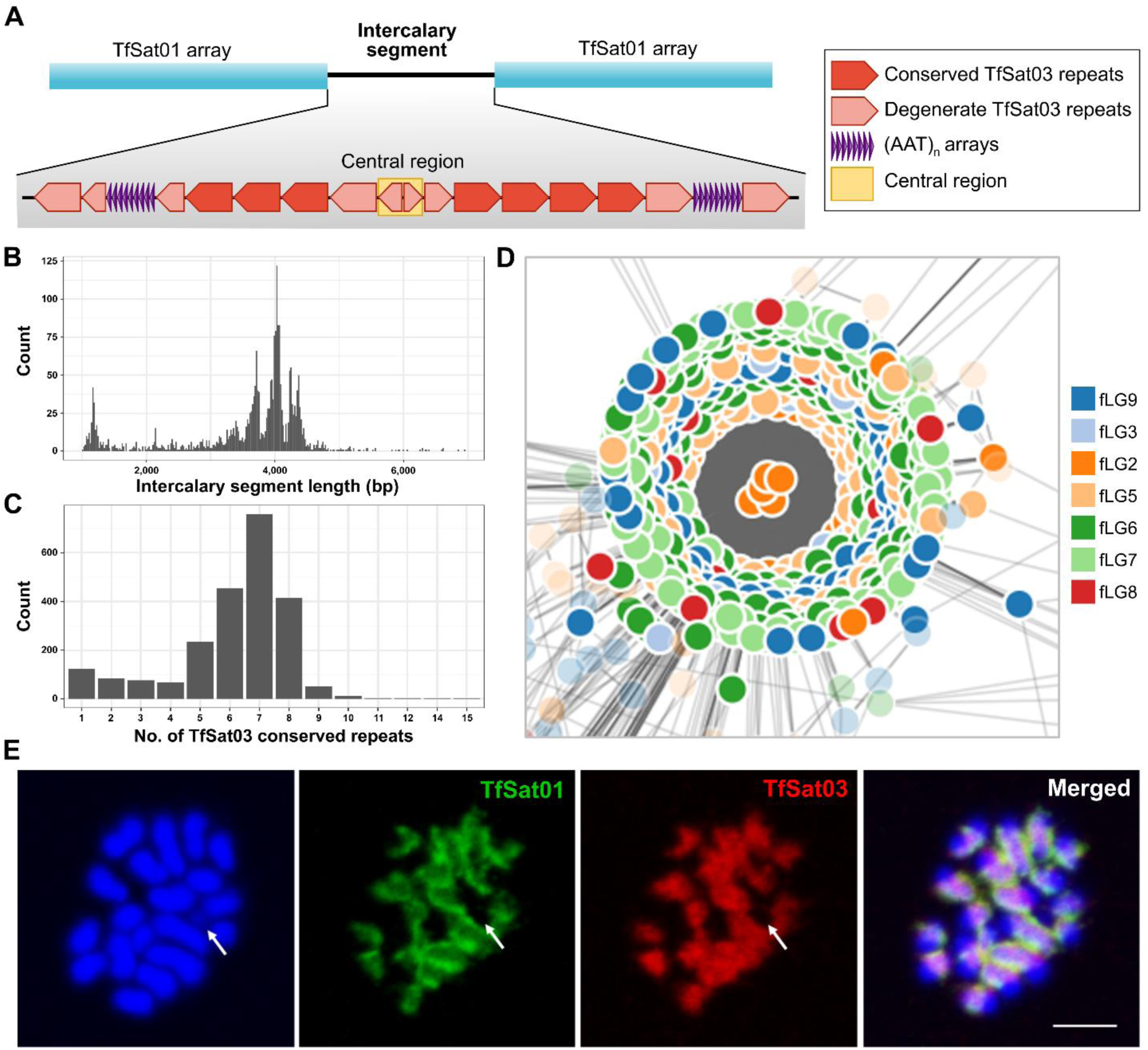
Organization of TfSat03 satDNA repeats in the intercalary segments between TfSat01 arrays. **A**) Schematic representation of the intercalary segments containing conserved and degenerate TfSat03 repeats and (AAT)_n_ microsatellite arrays. The yellow rectangle marks the central region where the degenerate copies of TfSat03 change orientation. **B**) The distribution of intercalary segments lengths containing TfSat03 repeats. **C**) The number of conserved TfSat03 complete repeats per segment in the analyzed intercalary segments. **D**) Graph networks of central region sequences based on their sequence similarity relationship. The figure shows a detail of the graph network (the entire graph network can be found in Supplementary Figure S10) and highlights a cluster consisting of 100% identical sequences originating from different chromosomes. Each dot represents one sequence, and the color of the dot indicates the chromosome on which the sequence is located. The legend of the color-coded chromosomes is shown next to the graph. **E**) Co-localization of TfSat01 (green) and TfSat03 (red) satDNAs on the *T. freemani* male metaphase chromosomes stained in DAPI (blue). An arrow points to the y_p_ chromosome lacking the TfSat01 and TfSat03 signals. The bar represents 3 µm.

A ∼110 bp long central region (marked by the yellow rectangle in **Fig. 3A**), where the degenerate TfSat03 repeats change orientation, represents the midpoint of the dyad symmetry, potentially resulting in a 13 bp loop (**Suppl. Fig. S9**). Interestingly, the sequence of the central region is so degenerate that the partial similarity to TfSat03 is barely recognizable. Despite its degeneracy compared to TfSat03, the ∼110 bp central region is highly conserved across intercalary segments throughout the genome, showing an average pairwise identity of 97.1% (**Suppl. Data S2**). We used undirected graph networks generated from distance matrices to visualize relationships between central regions from different intercalary segments. The graph networks revealed some small chromosome-specific clusters (**Suppl. Fig. S10, Suppl. Data S3**), but the largest cluster with over two hundred 100% identical sequences of the central region is formed from the intercalary segments distributed on different chromosomes (**Fig. 3D**). Thus, we reason that the ∼4 kb long dyad symmetries of TfSat03-based intercalary segments, along with their preserved central region, represent a conserved trait. In other words, our analysis shows that the centromeric regions of *T. freemani* are composed of alternating arrays of TfSat01 and TfSat03 satDNAs, with the arrays of both satellites exhibiting a strong tendency towards macro-dyad symmetries.

To further corroborate the relationship between these two satDNAs *in situ*, we performed double-color FISH. This confirmed the colocalization of TfSat01 and TfSat03 in broad regions of the 19 *T. freemani* chromosomes in males (**Fig. 3E**). Notably, neither TfSat01 nor TfSat03 signals were detected on the y_p_ chromosome (**Fig. 3E**, marked by an arrow). The fact that the y_p_ chromosome lacks the two most abundant and widespread *T. freemani* satDNAs opens the question of y_p_ satellite profile.

### 4. TfSat04, the male sex chromosome specific satDNA

As mentioned, some *T. freemani* satDNAs show mutual similarities in their nucleotide sequences. TfSat03 is one of them, belonging to the SF1 superfamily, that also comprises satDNA TfSat04 (**Suppl. Table S3**). In contrast to TfSat03, TfSat04 is a low-copy-number satellite that makes up only 0.07% of the genome. Beyond the 80-fold difference in genome abundance, the two satDNAs also differ in the structure of their repeat unit. The monomer TfSat03 is a 340 bp HOR whose three subunits A, B and C are 113-114 bp long and share pairwise similarities of 65.8-73.0% (**Suppl. Fig. S2A**), while TfSat04 is based on a 112 bp long repeat that corresponds to TfSat03 subunits sharing with them 65.8-72.6% similarity (**Fig. 4A, Suppl. Fig. S1A**). The principal component analysis (PCA) of all 159 TfSat04 monomers mapped in the Tfree1.0 assembly and 300 randomly subsampled TfSat03 subunits clearly separated the sequences into four distinct clusters, reflecting their provenience (**Fig. 4B**). According to the PCA, TfSat04 monomers are most closely related to TfSat03 subunit A.

**Figure 4.**
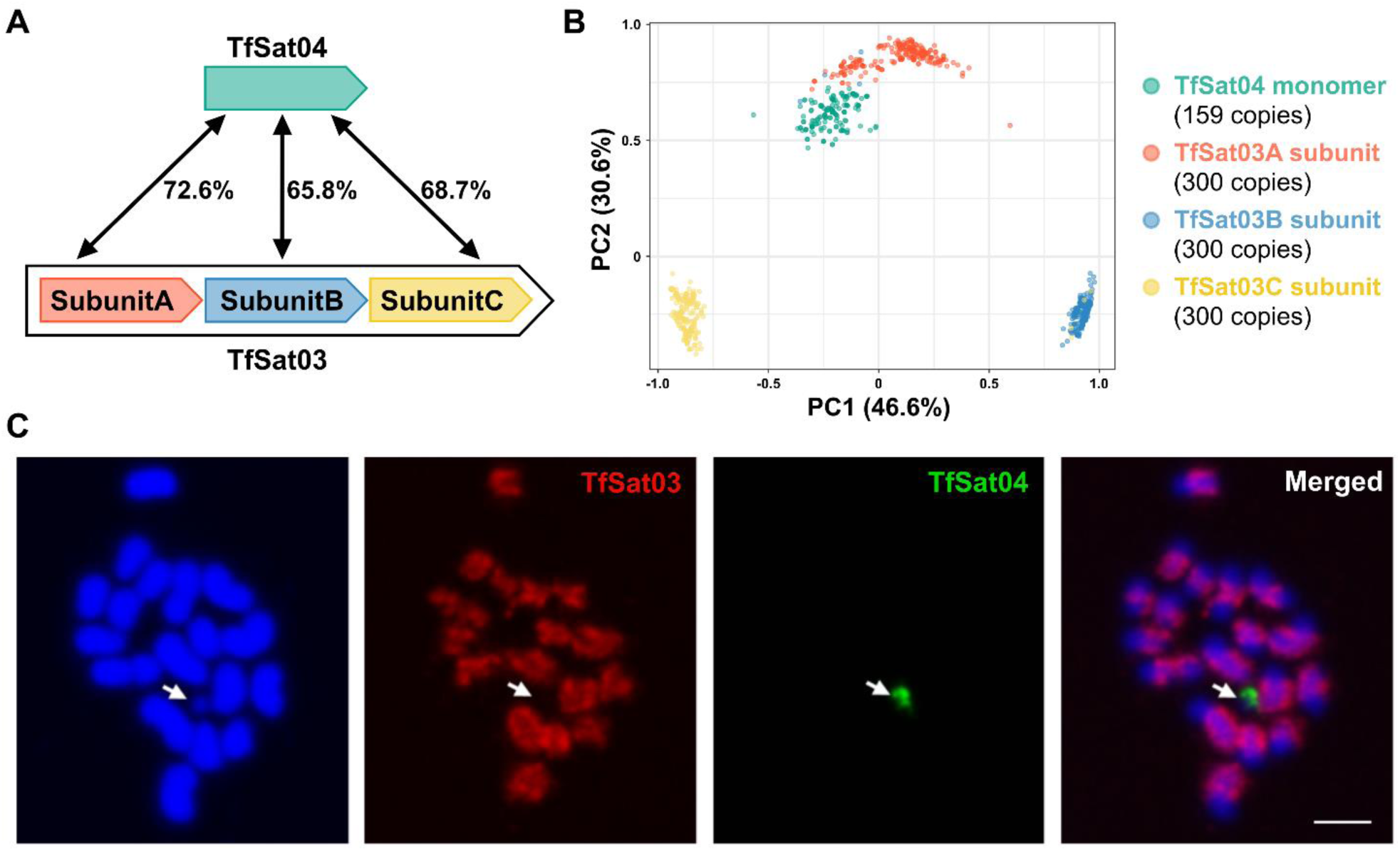
Relationships between TfSat04 and TfSat03 satDNAs. **A**) Schematic representation of the differences in the structure of the basic repeat unit and the pairwise similarities between the TfSat04 monomer and the three subunits of TfSat03. **B**) PCA clustering of 159 TfSat04 monomer copies and TfSat03 subunits A, B and C extracted from the 300 randomly selected TfSat03 repeats. The monomers and subunits are represented by color-coded dots. The color-coded legend is provided next to the PCA plot, and the proportions of variance for the first two principal components, PC1 and PC2, are indicated on the axes in parentheses. **C**) Co-localization of TfSat03 (red) and TfSat04 (green) satDNAs on the *T. freemani* male metaphase chromosomes stained in DAPI (blue). The arrow points to the y_p_ chromosome, on which there is no TfSat03 signal, but a TfSat04 signal that is exclusively present on this chromosome. The bar represents 3 µm.

Since the Tfree1.0 assembly contains only a single continuous array of TfSat04 annotated on chromosome fLG7, by performing FISH we expected to detect it on one chromosome. Indeed, only one TfSat04 signal was obtained on chromosome spreads, but to our surprise the signal was located on the y_p_ chromosome. Double-color FISH confirmed that the positions of TfSat03 and TfSat04 are mutually exclusive, i.e. while TfSat03 is localized at 19 chromosomes and not on y_p_, TfSat04 is exclusively present on y_p_ (**Fig. 4C**). FISH on female chromosome spreads, where no TfSat04 signal was detected, confirmed that TfSat04 satDNA is specific for the male sex chromosome.

The experimental detection of TfSat04 on the y_p_ chromosome prompted us, as the authors of the Tfree1.0 genome assembly, to reexamine the original contigs that made up Tfree1.0. After revision (explained in detail in Material and Methods section), we concluded that the TfSat04-containing contig, located at the end of chromosome fLG7 in the Tfree1.0 assembly, belongs to the male chromosome y_p_. Therefore, following this finding, we will curate the Tfree1.0 assembly by upgrading it to a new version Tfree1.1, in which the 2.2 Mb long end of chromosome fLG7 is separated and proposed as the y_p_ chromosome.

### 5. Orthologous satDNAs between *T. freemani* and *T. castaneum*

To explore the evolutionary trends in the satellitome of *T. freemani*, we included the orthologous satDNAs of the most closely related species, *T. castaneum*, in the study.

#### The origin of the centromeric major satDNAs of the sibling species

The siblings *T. freemani* and *T. castaneum* differ by completely unrelated, centromerically located major satDNAs, TfSat01 and TCAST. Therefore, we first addressed the origin of these satellites.

By analyzing 228,060 TfSat01 monomer copies annotated in the Tfree1.0 assembly, we found that 93.3% of them exhibit sequence similarity to the consensus greater than 90% (**Suppl. Fig. S5B**), indicating that TfSat01 is a highly homogeneous satellite. Nevertheless, the histogram of the similarity distribution also revealed a distal peak corresponding to 82% similarity (indicated by the black arrow in **Suppl. Fig. S5B**). We found that these lower-similarity copies do not belong to tandemly organized TfSat01 repeats, but are integrated parts of the 1106 bp long repeat units of *T. freemani* satDNA TfSat02 (**Fig. 5A, Suppl Fig. S11**). Interestingly, TfSat02 repeats, which constitute 1.6% of the genome (**Fig. 1A**, **Suppl. Table S3**), are located outside the TfSat01 areas, often far from centromeric regions (**Fig. 5B**). This raised the question of the evolutionary relationship between the two satellites, i.e., which satDNA preceded the other: 1) was TfSat01 an ancestral satDNA whose monomeric unit invaded the TfSat02 precursor sequence, or 2) was a fragment of the TfSat02 sequence excised and amplified into tandem arrays that gave rise to the present major satDNA TfSat01. To answer this, we searched the genome of the sibling species *T. castaneum* for sequences that might be related to TfSat01 or TfSat02. We did not detect any tandemized TfSat01-like copies in *T. castaneum*. However, we discovered that TfSat02 is the ortholog of the previously described *T. castaneum* satDNA TCsat15 (Gržan et al. 2023), sharing a full monomer length of 1106 bp and 85.2% sequence similarity (**Suppl. Fig. S11**). Like TfSat02, the TCsat15 repeat contains a 166 bp segment corresponding to TfSat01 monomer (**Fig. 5A**, **Suppl. Fig. S11**), which implies that the TfSat02/TCsat15 predecessor sequence contained the 166 bp TfSat01 precursor. From this we conclude that the major satellite TfSat01 is evolutionarily younger and that it emerged from TfSat02. A comparison of the TfSat01 consensus sequence with the corresponding 166 bp segments of TfSat02 and TCsat15 revealed that the TfSat02 and TCsat15 segments are more similar to each other than to the TfSat01 consensus (**Fig. 5A**). In addition to the simple consensus comparison, we conducted a more comprehensive PCA, which included 10,000 randomly subsampled TfSat01 monomers, all TfSat02 166 bp segments annotated in the *T. freemani* genome assembly Tfree1.0, and all TCsat15 166 bp segments annotated in the *T. castaneum* genome assembly TcasONT. PCA strongly grouped the TfSat01 monomers into a separate, distant cluster (**Fig. 5C**). In line with the fact that no tandemized TfSat01-like copies were mapped in *T. castaneum*, we conclude that TfSat01 was not present as canonical satDNA in the common ancestor, but instead is a trait of the *T. freemani* genome.

**Figure 5.**
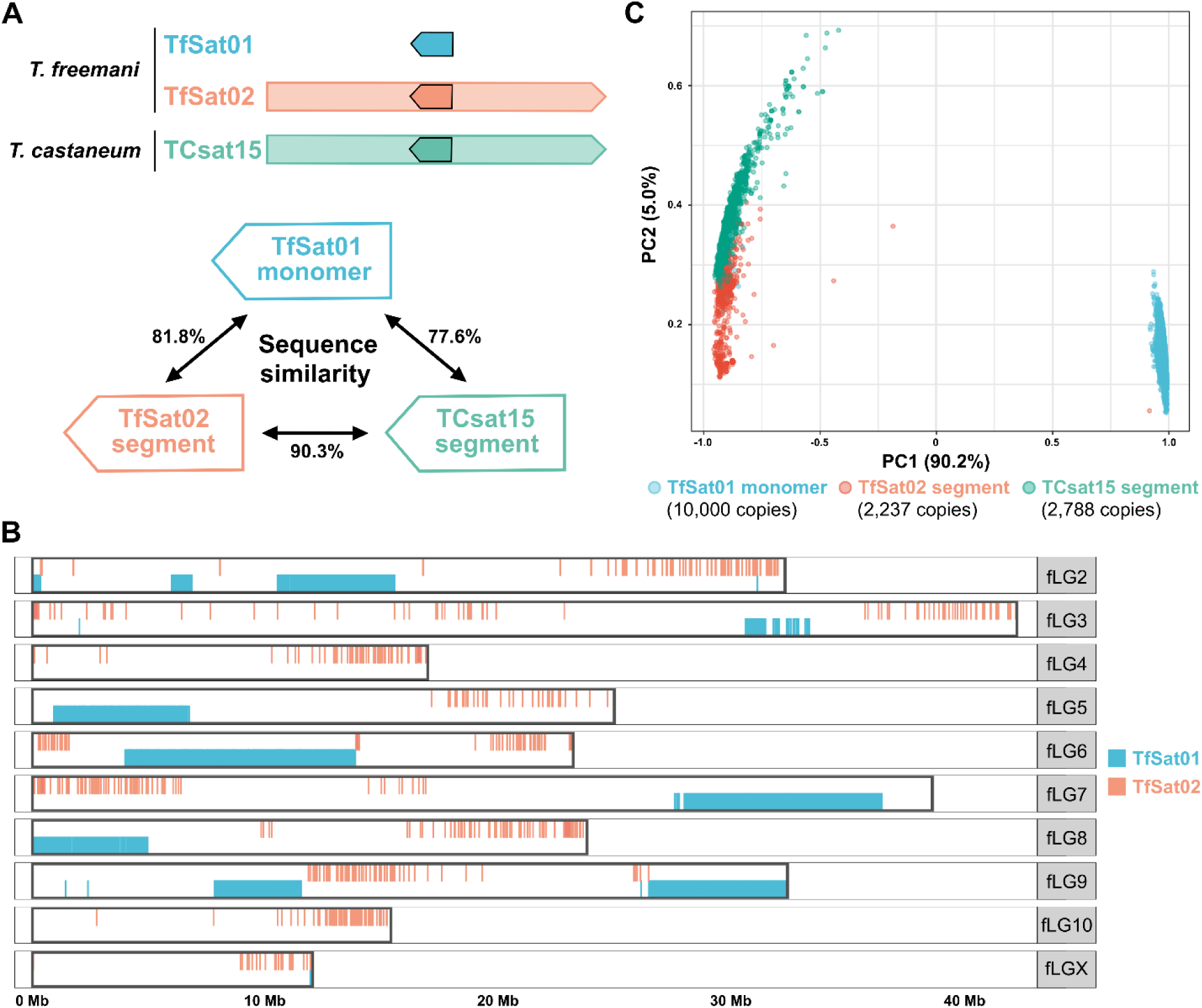
Relationship between *T. freemani* satDNAs TfSat01 and TfSat02, and *T. castaneum* satDNA TCsat15. **A)** Schematic representation of the TfSat01, TfSat02, and TCsat15 repeat units. The blue arrow represents the TfSat01 monomer, while the red and green arrows within the TfSat02 and TCsat15 units, respectively, indicate the position of the 166 bp segments that correspond to the TfSat01 monomer sequence. The percentages of nucleotide sequence similarity between the consensus sequences of TfSat01 monomers and the corresponding 166 bp segments of TfSat02 and TCsat15 are indicated. **B)** Distribution of satDNA TfSat01 and TfSat02 repeats mapped at the Tfree1.0 genome assembly along the *T. freemani* chromosomes fLG2-fLGX. **C)** PCA clustering of TfSat01 monomers and 166 bp segments from TfSat02 and TCsat15. The analysis included 10,000 randomly selected TfSat01 monomers and 2,237 copies of TfSat02 segments detected in the *T. freemani* genome assembly Tfree1.0, and 2,788 copies of TCsat15 segments annotated in the *T. castaneum* genome assembly TcasONT. Each dot represents a sequence of a single monomer/segment, and the color-coded legend is provided below the PCA plot. The proportion of variance for the first two principal components, PC1 and PC2, is shown on the axes in parentheses.

To decipher the origin of the *T. castaneum* major satDNA TCAST, we searched for its possible presence in *T. freemani*. However, we did not detect any TCAST tandemized copies in the Tfree1.0 assembly. Instead, we detected a TCAST transposon-like element that has been previously reported in the *T. castaneum* genome (Brajković et al. 2012). Resembling to a DNA transposon, the TCAST transposon-like element is 1093 bp long and contains two segments corresponding with 83% similarity to a TCAST monomer and its truncated version in reverse orientation, partially overlapping with ∼290 bp inverted termini (**Suppl. Fig. S12A**). We found this transposon-like element scattered on all chromosomes in both species, except for the *T. castaneum* X chromosome (**Suppl. Fig. S12B**). When compared, the copies of the TCAST transposon-like elements from *T. castaneum* and *T. freemani* generally separate in species-specific groups (**Suppl. Fig. S12C**). Considering the 80% sequence similarity and wide genomic distribution in both species, we hypothesize that the TCAST transposon-like element was present in the ancestral genome, and served as a source from which the 360-bp TCAST satDNA, now accounting for 17% of the *T. castaneum* genome, was derived.

#### The orthologous centromeric minor satDNAs

TfSat03 is the second most abundant satellite in *T. freemani*, so we next searched for its ortholog in *T. castaneum*. It turned out to be Cast7, one of the most variable *T. castaneum* satDNA with an estimated genomic proportion of 0.2% (Pavlek et al. 2015; Volarić et al. 2024). Cast7 is based on ∼109-114 bp repeat units, which correspond to the TfSat03 subunits and show on average 67.2-79.3% similarity to them (**Suppl. Fig. S13**). Even more fascinating than the nucleotide similarity between Cast7 and TfSat03 is their orthologous organization characterized by intermingling with the major satDNAs and a propensity for dyad symmetry (**Fig. 6A**). Namely, in *T. castaneum*, Cast7 arrays are primarily located between the TCAST arrays in the (peri)centromeric regions (Volarić et al. 2024). Notably, Cast7 also exhibits changes in the orientation of its copies within the arrays, and we discovered that the major satellite TCAST does the same. While the TCAST-Cast7 arrangement in *T. castaneum* is more irregular than the juxtaposition of TfSat01 and TfSat03 in *T. freemani*, the positional and organizational orthology of the minor satellites Cast7 and TfSat03 is evident despite the fact that the major satellites, TCAST and TfSat01, are different (**Fig. 6A**).

**Figure 6.**
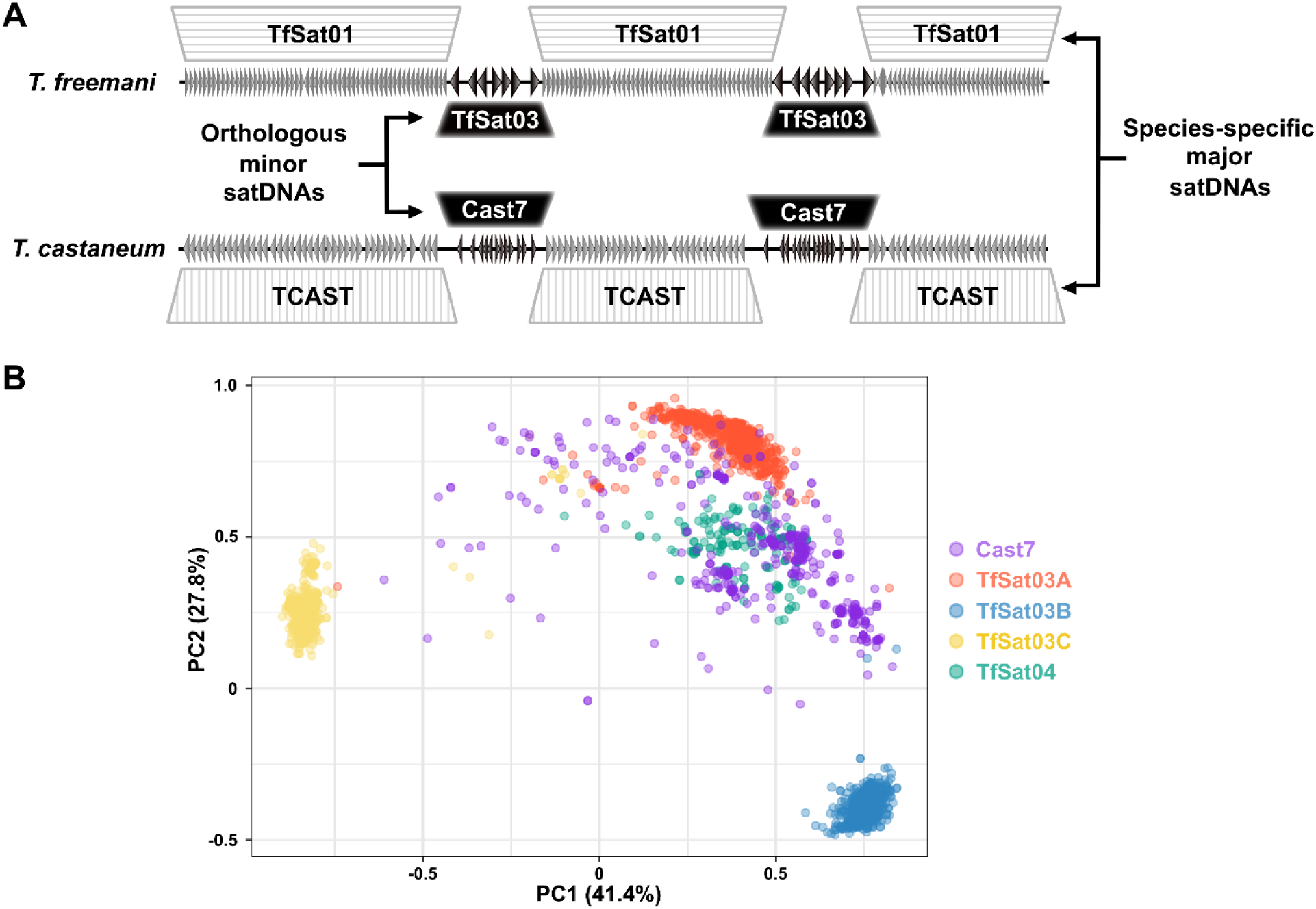
The comparison of satDNAs in the centromeric regions of the sibling species *T. freemani* and *T. castaneum*. **A)** A schematic illustrating the long-range organization of the species-specific major satDNAs TfSat01 and TCAST, intermingled with orthologous minor satDNAs TfSat03 and Cast7, respectively, in the centromeric regions of *T. freemani* and *T. castaneum*. **B)** PCA clustering of the repeats of the orthologous satDNAs Cast7, TfSat03 and TfSat04. The analysis included 1000 randomly selected Cast7 monomers from the TcasONT assembly, together with subunits A, B, and C from 1000 randomly selected TfSat03 repeats and all 159 TfSat04 monomers annotated in Tfree1.0 assembly. The sequences are represented by dots according to the color-coded legend provided next to the PCA plot.

Given the relatedness of the satellites TfSat03 and y_p_-specific TfSat04, we further investigated the relationship between Cast7, TfSat04 and TfSat03 subunits by analyzing randomly selected individual monomer/subunit sequences. According to PCA (**Fig. 6B**), Cast7 monomers intermingle between clusters of TfSat04 monomers and TfSat03_subunit A, confirming the observed highest similarity between their consensuses (**Suppl. Fig. S13**). The increased connection between Cast7-TfSat04-TfSat03_subunitA sequences could be an indication of ancestral sequence preservation, but potentially also a sign of a certain functionality.

#### The orthologous low-copy-number satDNAs

In the search for orthologous satellites, besides TfSat02, TfSat03 and TfSat04, we identified 11 additional *T. freemani* satDNAs with orthologs in the previously characterized *T. castaneum* satellites (**Suppl. Table S3)**. The alignments of the species consensuses show pairwise similarities from 63.8% to 94.8% (**Suppl. Fig. S14**). To further explore the relationships between the monomeric copies of orthologous satellites, we annotated the orthologous repeats in Tfree1.0 and TcasONT and compared their sequences (**Suppl. Data S4**). The resulting PCA plots showed one of three patterns: 1) species-specific clustering (TfSat08, TfSat11, TfSat15, TfSat25) (**Fig. 7A**), 2) extensive mixing of repeats from both species (TfSat05, TfSat22, TfSat111) (**Fig. 7B**), 3) segregation of repeats from one species into multiple cluster (TfSat07, TfSat27, TfSat42, TfSat76) (**Fig. 7C**). We hypothesized that some of the remaining 120 *T. freemani* satDNAs might have orthologous copies that are not defined as satDNA in *T. castaneum*. This could be due to the genuine absence of their copies in *T. castaneum* or the incompleteness of the repetitive DNA regions in the *T. castaneum* Tcas5.2 assembly, which was used to define the *T. castaneum* satellitome (Gržan et al. 2023). To test this hypothesis, we searched the *T. castaneum* TcasONT assembly using all *T. freemani* satDNAs as queries, and we indeed identified additional 45 satellites recognizing related copies in *T. castaneum*. Based on the alignments of these individual copies (**Suppl. Data S4**), the PCA (**Suppl. Data S5**) revealed that these 45 orthologs also follow one of the three previously described patterns.

**Figure 7.**
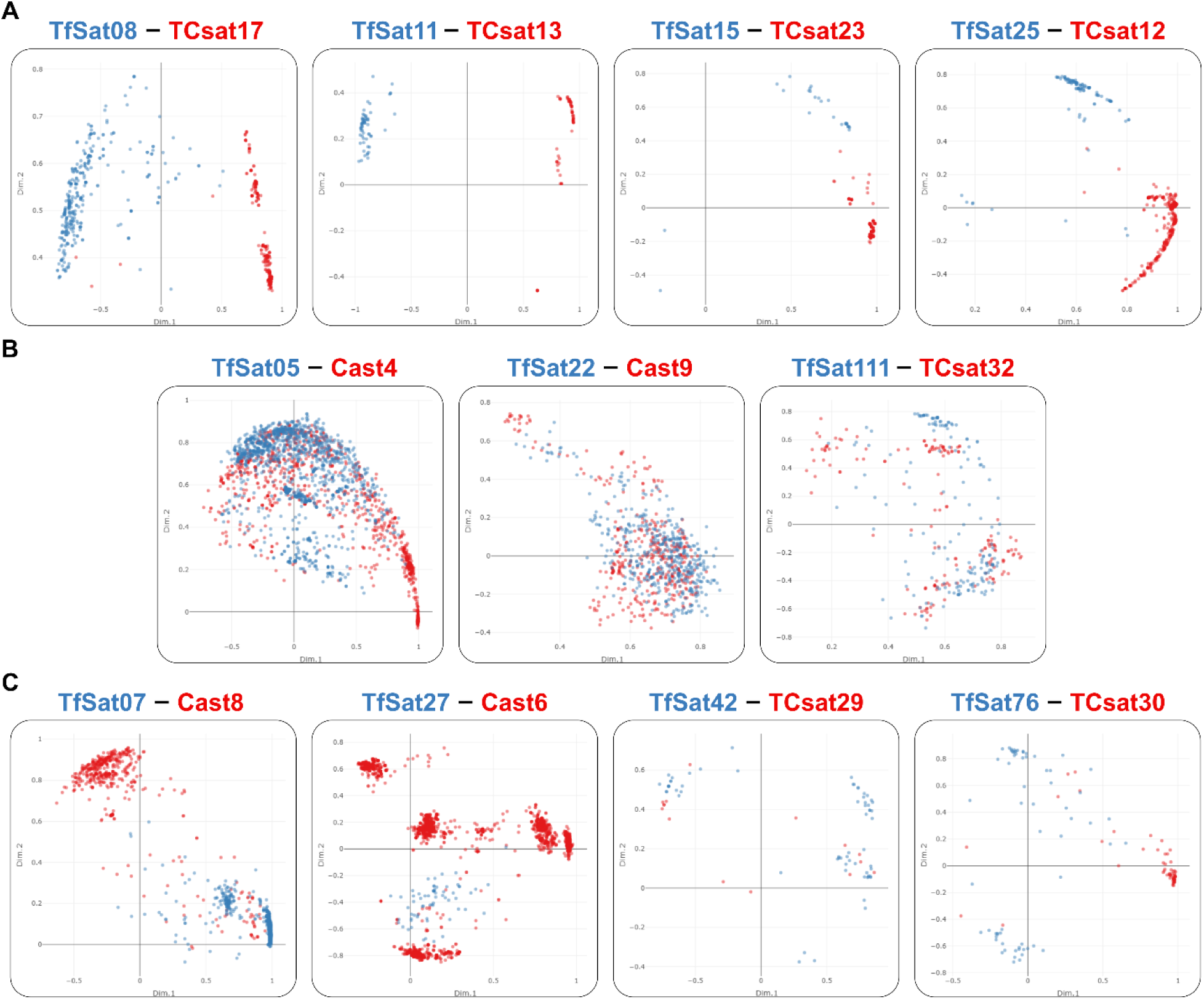
PCA plots of the *T. freemani* satDNAs (TfSat) repeats and their orthologs among known *T. castaneum* satDNAs (Cast, TCsat)showing: A) species-specific clustering, B) extensive interspecific mixing, C) segregation of repeats from one species in more than one cluster. The names of the orthologous satDNAs are indicated above each plot. The dots, colored according to the species of origin (*T. freemani* in blue, *T. castaneum* in red), represent monomer copies annotated in the *T. freemani* Tfree1.0 assembly and the *T. castaneum* TcasONT assembly. The PCA plots are based on the alignments shown in Supplementary Data S4. The interactive versions of the plots can be found in Supplementary Data S5.

### 6. Mechanisms of satDNA propagation

Being among the most variable sequences of eukaryotic genomes, satDNAs are subject to a very dynamic turnover. To infer possible mechanisms that have formed and probably actively remodel the satellitome of *T. freemani*, we first focused on the two most abundant satellites, TfSat01 and TfSat03. The intermingling of these two satellites across all chromosomes, with the exception of the sex chromosome y_p_, indicates that the spread of TfSat01 and TfSat03 is closely linked. In addition, the inversion points within TfSat01 and TfSat03 arrays, distributed in a highly conserved form at non-homologous chromosomes, suggest mechanisms that spread the two satellites together to different chromosomes. One potential mechanism for this collinear spread could be the expansion via extrachromosomal circular DNA molecules (eccDNAs) that contain the TfSat01/TfSat03 arrays and spread them by the reintegration into non-homologous chromosomes. To test the presence of TfSat01 and TfSat03 repeats in the eccDNA fraction, we employed two-dimensional (2D) agarose gel electrophoresis of the total genomic DNA. Subsequent Southern blot hybridizations with TfSat01 and TfSat03 specific probes showed, as expected, TfSat01 and TfSat03 signals corresponding to linear, chromosomal DNA, but also revealed their concomitance at the arc signals indicative of eccDNA, confirming that these two satDNAs indeed occur in the form of extrachromosomal circular molecules (**Fig. 8A**). Another way of spreading and exchanging these satellites between non-homologous chromosomes could be achieved by 3D interactions between different chromosomes. Indeed, in meiotic prophase I we occasionally observed the bouquet-like formations of non-homologous chromosomes huddled together by (peri)centromeric heterochromatin. By double-color FISH, we confirmed that TfSat01 and TfSat03 signals colocalize at these close associations of the (peri)centromeric regions (**Fig. 8B**). We conclude thus that the concomitance of the two satellites at multi-megabase long regions might be a consequence of the joint action of both mechanisms, (peri)centromeric 3D interactions at meiotic bouquet-like configurations and reintegration of eccDNAs.

**Figure 8.**
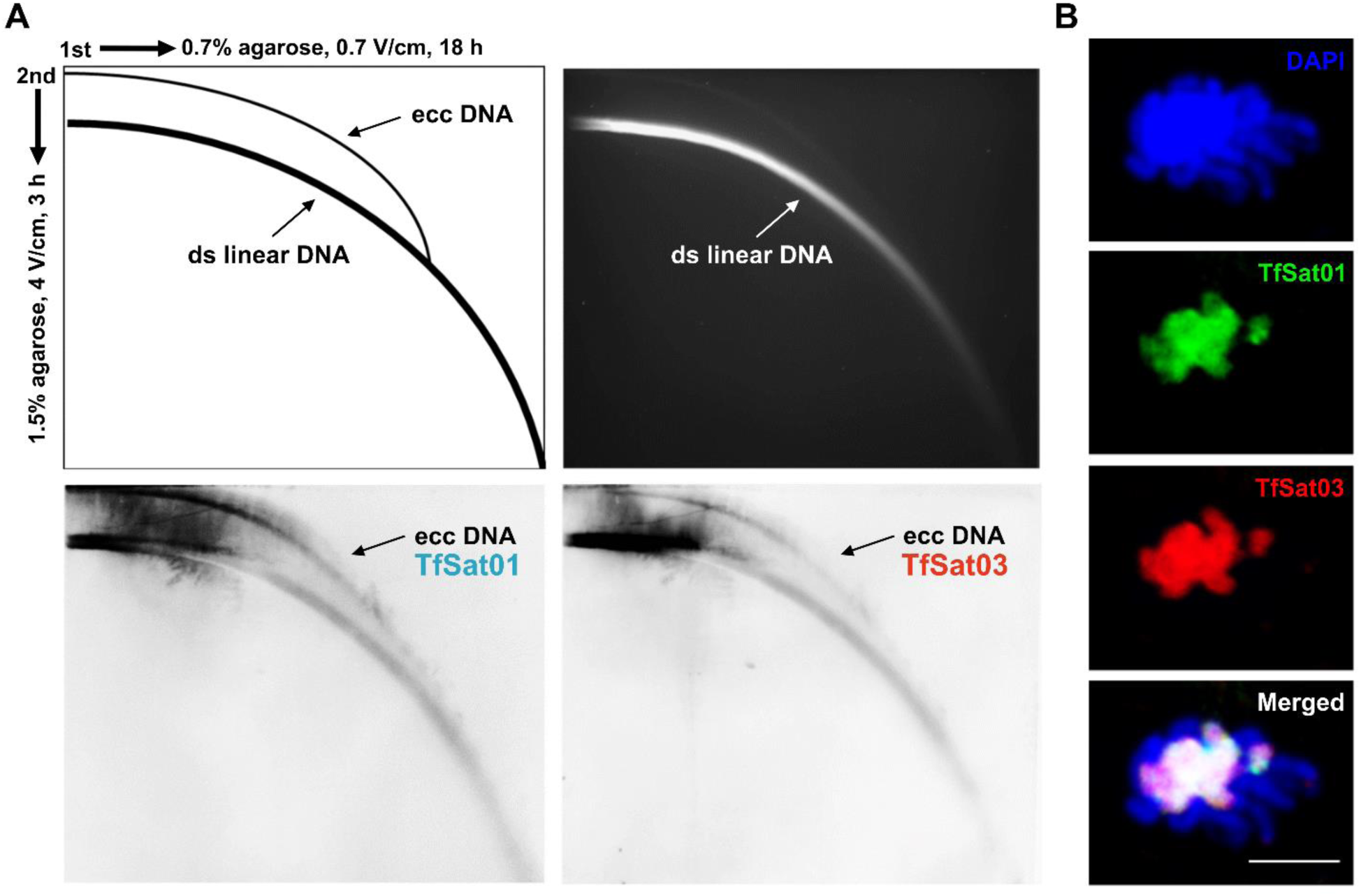
Experimental evidence of the presence of the TfSat01 and TfSat03 satDNA repeats in the extrachromosomal circular DNA (eccDNA) molecules and in prophase I associations of non-homologous chromosomes. **A**) Two-dimensional gel electrophoresis analysis of the eccDNAs in *T. freemani*. The schematic at the top left illustrates 2D gel electrophoresis and the migration patterns of linear and circular DNA forms. The ethidium bromide stained gel (the upper right panel) with 2D-electrophoretically separated *T. freemani* genomic DNA was Southern-blotted and hybridized with the TfSat01 probe (lower left panel) and the TfSat03 probe (lower right panel). Both blots show the signals corresponding to eccDNAs (indicated by the arrows). **B**) Meiotic bouquet-like formations of *T. freemani* chromosomes with joint heterochromatic regions of non-homologous chromosomes. Chromosomes (stained with DAPI, top panel) were subjected to double-color FISH analysis using specific probes for TfSat01 (green) and TfSat03 (red). The bottom panel shows the overlap of the two satDNAs. The bar represents 5 µm.

In addition to these two satellites, we observed potential mechanisms of propagation and expansion in other satDNAs. Partial sequence similarities of certain low-copy-number satDNAs with TEs (**Suppl. Table S3**) suggest that transposons may be a source of some tandem repeats in *T. freemani* and that transposition may be a mechanism contributing to their propagation. In the satDNA-TE linkage analysis, we found the orthologous satDNAs to be particularly informative. The clearest examples of association with TEs come from satDNAs related to the Rehavkus superfamily of DNA transposons, characterized by long terminal inverted repeats and a variable number of subterminal tandem repeats. Here we highlight the three cases. The first involves the satDNA TfSat15, which, like its ortholog TCsat23 in *T. castaneum*, forms short tandem arrays of several consecutive monomers in the inverted termini of the Rehavkus-1_TC transposon (**Suppl. Fig. 15A**). Analysis of the orthologs showed that the copies of TfSat15 and TCsat23 cluster into species-specific groups (**Fig. 7**), a pattern also observed for the rest of the transposon sequence (**Suppl. Fig. 15A**). A second example is the satellite TfSat23, which resides in the inverted termini of the Rehavkus-3_TC transposon. The repeats of TfSat23 and its orthologous copies in *T. castaneum* also evolve according to the principles of concerted evolution, as does the central part of the transposon sequence (**Suppl. Fig. 15B**). However, Rehavkus-3_TC transposons in *T. freemani* typically contain 6-8 TfSat23 repeats in their inverted termini, while the number of satellite repeats in transposon copies in *T. castaneum* is generally larger, reaching up to 50 consecutive monomers. The third and most illustrative example of how transposon-integrated repeats can expand remarkably into long tandem arrays is TfSat25. In *T. freemani*, this repeat makes up 2-12 consecutive copies in the inverted termini of the Rehavkus-1_TC-like elements. In *T. castaneum*, on the other hand, its orthologous satellite TCsat12 forms very long stretches, with the longest reaching 80.5 kb and comprising 522 consecutive repeats on chromosome LG4 (**Suppl. Fig. 15C**). Considering that TfSat25 was found in the *T. freemani* assembly exclusively in association with Rehavkus-like elements, we conclude that the intense expansion of this repeat in *T. castaneum* occurred after the separation of the siblings. Although TfSat25 and TCsat12 still show a high similarity of 94.8% in nucleotide sequence (**Suppl. Fig. S14**), it is evident from the clustering of their repeats (**Suppl. Fig. 15C**) that they have evolved in a species-specific direction after the species split.

We also noted the effect of concerted evolution on satDNAs that are not linked to TEs. For example, the orthologous satDNAs TfSat08 and TCsat17 were found in both species on chromosome (f)LG3 in an array with a similar number of copies (339 in *T. freemani*, 290 in *T. castaneum*). Synteny analysis revealed that the TfSat08/TCsat17 satellite block is embedded in the same environment in both species, with 90.3% and 92.1% sequence similarity in the 10 kb flanking regions (**Suppl. Fig. 16**). PCA analysis, however, showed a clear species-specific grouping of individual TfSat08 and TCsat17 repeats (**Fig. 7A**). This indicates that the satDNA was already present in the ancestral species in an array of about 40 kb in length, but after the separation of the species, sequence conversion formed its species-specific variants.

Some satDNAs, such as TfSat06, show intense interchromosomal homogenization, as evidenced by their distribution across all chromosomes of *T. freemani* without chromosome-specific clustering (**Suppl. Fig. S17A**). Interestingly, TfSat06 arrays occur in an organizational form reminiscent of long, sometimes over 11 kb long, inverted termini of transposon-like elements, which show the greatest partial similarity to Rehavkus-1_TC sequence (**Suppl. Fig. S17B**), so we assume that transposition is responsible for the spreading of TfSat06. Since we identified only one orthologous 20-repeat array in *T. castaneum*, we assume that the expansion and interchromosomal homogenization of this satellite in *T. freemani* occurred after the species split. On the other hand, the orthologous satDNA pair TfSat22-Cast9 is evidence for a satellite that probably began its homogenization between different chromosomes in the ancestral genome. Namely, we saw a strong admixture of orthologous copies from the two species, but we also detected a strong interchromosomal admixture when we analyzed repeats within a species (**Suppl. Fig. S18**).

Finally, for some low-copy satDNAs, intraarray homogenization was observed. For example, the satDNA TfSat54, located on chromosome fLG10, forms two short arrays with repeats grouped in separate clusters (**Suppl. Fig. S19A**). Similarly, the satDNA TfSat97 shows an array-specific clustering of repeats from two arrays on chromosome fLG3 (**Suppl. Fig. S19B**). In both cases, however, the arrays of these very low-copy-number satDNAs do not exceed 20 tandem repeats, which raises the question of whether these are just satDNA “seeds” that may either spread or eventually disappear in the future.

## DISSCUSSION

In this study, we identified 135 satDNAs comprising 38.7% of the flour beetle *T. freemani* genome, one of the most satellite-rich genomes described so far. Recently, species with even larger numbers of satDNAs have been reported, such as the frog *Proceratophrys boiei* with 226 satellites (João Da Silva et al. 2023) or the crayfish *Pontastacus leptodactylus* with 258 satDNAs (Boštjančić et al. 2021), but their genomes are 10-60 times larger than *Tribolium*’s. While satDNA cataloging provides general information about the satellitome content, studying the long-range organization and structure of satDNA arrays is essential for understanding the evolutionary dynamics of these sequences, which was the ultimate goal of our work. To achieve this, we took advantage of: 1) the high-quality *T. freemani* reference genome (Volarić et al. 2022) based on PacBio HiFi reads, and 2) the close relationship of *T. freemani* with the coleopteran model *T. castaneum*, which also has a high-quality assembly (Volarić et al. 2024) and a well-defined satellitome (Ugarković, Podnar, et al. 1996; Pavlek et al. 2015; Gržan et al. 2023). These two points enabled us to discover orthologous satellites and investigate satDNA dynamics in the extremely satDNA-abundant genomes, with particular attention to centromeric satellites.

### The evolutionary aspects of Tribolium satellitomes

The origin of satDNAs, especially those that dominate genomes, is an intriguing but rarely answered question, despite numerous satDNA studies. Here, we revealed the origin of the genome-prevailing and species-specific centromeric major satellites in the siblings *T. freemani* and *T. castaneum*. We discovered that the *T. freemani* major satDNA TfSat01, which makes up one third of the *T. freemani* genome, arose from a sequence segment of another satellite that is still present in both genomes. Apart from the fact that the primordial satellite continues to exist in the current genomes of the siblings, an equally intriguing question is why only 15% of its sequence length has evolved into the new satellites. The proliferation of a dominant satDNA from a segment of a repetitive sequence is also demonstrated by the *T. castaneum* major centromeric satDNA TCAST. Namely, a DNA transposon-like element harboring TCAST was found scattered in the genome of *T. castaneum* (Brajković et al. 2012), but we also detected the same DNA transposon-like element in the *T. freemani* genome. The previous work on TCAST-like elements in *T. castaneum* also disclosed CR1-3_TCa retrotransposon, that contains a segment of TCAST sequence (Brajković et al. 2012), and the authors pointed out the possibility that the TCAST satellite derived from this retrotransposon. However, we could not detect any CR1-3_TCa retrotransposon copies in *T. freemani*. Therefore, we believe that it is more likely that TCAST originated from the ancestrally present DNA transposon, while CR1-3_TCa retrotransposon in *T. castaneum* might captured part of the TCAST sequence by insertion into the TCAST satDNA array.

The *T. freemani* and *T. castaneum* major satDNAs apparently evolved from different types of repetitive sequences, TfSat01 from a satellite and TCAST from a DNA transposon. The common link between these source elements is their genome-wide distribution and presence on all autosomal chromosomes. We conclude that the ancestral sequence TfSat02/TCsat15, which gave rise to satellite TfSat01, must have been widely scattered in the ancestral genome, because in today’s genomes both widespread satellites, TfSat02 and TCsat15, often occur in shorter segments with one or a few repeats (this work, (Gržan et al. 2023)), reminiscent of a dispersed element. We therefore hypothesize that the widespread distribution of an element throughout the genome is, if not a prerequisite, at least a desirable property that sequences with the potential to be the source of satDNAs exhibit. Under circumstances of intense DNA turnover, for example in some stress situations, dispersed elements might have a greater potential to distribute and exchange sequences with their counterparts from distant regions and/or non-homologous chromosomes through mechanisms that drive satDNA turnover. Importantly, we found neither tandemized copies of the *T. freemani* major satellite in the *T. castaneum* genome, nor tandemized copies of the *T. castaneum* major satellite in *T. freemani*. Therefore, we conclude that the proliferation of the major satDNAs in *T. freemani* and *T. castaneum* occurred simultaneously and possibly rapidly. The striking amplification and propagation of a sequence from the ancestor’s genome into a dominant satDNA of the offspring genome could be explained by the satDNA library theory (Fry and Salser 1977), which has been experimentally confirmed in different organisms (Meštrović et al. 1998; Silva et al. 2017). However, the satDNA library hypothesis assumes the propagation of the entire pre-existing repeat, whereas the relationship between TfSat01 and TfSat02 in *T. freemani* shows that it is possible that only a segment of a satellite monomer proliferates into a novel and highly abundant satDNA. This discovery expands the current understanding of the satDNA library concept, and if more such examples are found in other organisms, this could eventually help to identify the features that qualify a sequence (or its segment) to be propagated in the most copious satellite in a genome.

The low-copy-number satDNAs, whose number prevail in the *T. freemani* satellitome, show evolutionary statuses from species-specific to orthologous between the two siblings. In both groups there is a large number of satellites that show partial similarities with transposable elements (TEs). Links between TEs and satDNAs have been observed for some time, and there is growing evidence that TEs are a prolific source of satellite sequences (reviewed in Zattera and Bruschi 2022). In this respect, studies based on the long-range organization of repetitive sequences are particularly revealing. High-quality genome assemblies of the three species from the *Drosophila virilis* group showed that much of the divergence in genome composition between the sister species is due to Helitron TE-related tandem repeats (Flynn et al. 2024). Another TE-tandem repeat association in the *virilis* group involves satDNA repeats, which occur in large inverted termini of the foldback DNA transposon *Tetris* (Dias et al. 2014). Characterization of TE-satDNA associations in *T. freemani* highlighted Rehavkus, the foldback DNA transposons that, like *Tetris*, carry tandem repeats at their inverted ends. It has been shown that an increased number of tandem repeats in the inverted termini of foldback elements affects their transposition by strengthening transposase binding and increasing excision frequency (Liu and Wessler 2017), which may explain the benefit for foldback TEs to accommodate satellite repeats. Interestingly, orthologous satDNAs associated with Rehavkus elements in *Tribolium* change according to concerted evolution, as does the rest of the transposon sequence. While some repeats can remain cocooned in the inverted termini of TEs for a long time, some orthologs clearly demonstrated the amplification burst from transposon-residing repeats to an 80 kb long satellite array.

It is reasonable to assume that TEs serve not only as a source of satellite sequences, but also as a means of propagation. Transposition could be one of the efficient mechanisms by which satellite sequences spread throughout the genome and thus also in euchromatin regions that were dogmatically thought to be deprived of satDNAs. In *T. freemani*, we found most of low-copy-number satDNAs in the vicinity of genes and TEs, which is consistent with the results for the low-copy-number satDNAs in *T. castaneum* (Gržan et al. 2023). In *T. castaneum*, the extensive spread of even more abundant satDNAs in gene-rich regions was revealed (Volarić et al. 2024), following the example of dynamic euchromatic satDNAs described in *Drosophila melanogaster* and its three closest relatives (Sproul et al. 2020). A growing number of studies documenting the presence of satDNA arrays in euchromatic regions testify that euchromatin is not inhospitable to tandemly repetitive sequences (Cabral-de-Mello et al. 2023; Rico-Porras et al. 2024). However, functional analyses are required to clarify whether the spread of satDNAs in euchromatin is function-driven or whether these arrays are just tolerated scattered repetitive “seeds” with potential to expand.

### SatDNA-rich centromeric regions in Tribolium

Consistent with the satellite-rich genomes, the centromeric regions of *Tribolium* abound in satellites. In *T. castaneum*, satDNA-rich and unusually extended centromeres, so-called metapolycentromeres, comprise almost half the length of individual chromosomes (Gržan et al. 2020), and the FISH results in this work reveal a comparable situation in *T. freemani*. Such abundance brings into discussion the criteria that a sequence must fulfill to become a centromere-dominant satellite, but also the question of whether a sequence or a structure is more important. The long-range organization of satDNAs in the *T. freemani* (peri)centromeric regions pointed out dyad symmetry as the most conspicuous feature. Dyad symmetries within satellite monomers have the potential to form secondary structures such as hairpins or cruciforms, and enrichment with the predicted non-B-form DNA structures was found in the centromeric repeats of different animal and plant species (Kasinathan and Henikoff 2018; Liu et al. 2023). It has been suggested that the role of such non-canonical DNA structures in centromeric regions may be to facilitate the loading of centromere-specific nucleosomes (Talbert and Henikoff 2022). Short inverted repeats (<10 bp), which could potentially form non-B-form DNA structures, are present in the 166 bp long monomers of the *T. freemani* major satDNA TfSat01 (Juan et al. 1993), as well as in the 360 bp long monomers of the *T. castaneum* major satDNA TCAST (Ugarković, Podnar, et al. 1996). In addition to short dyad symmetries within monomers, here we revealed that *T. freemani* centromeric regions are populated with intermingled arrays of major TfSat01 and minor TfSat03 satDNA, both of which exhibit striking macro-dyad symmetries based on their several kb long inverted subarrays. The high frequency and preservation of these macro-dyad symmetries inevitably raises the question of what significance they might have. Since there is no evidence in the literature for stable hairpins or cruciforms formed by multiple kb long inverted sequences, we hypothesize that segments in which satellite subarrays/monomers change orientation and possibly form DNA loops could be key components of macro-dyad symmetries. The potential loops could contribute to the compaction of centromeric chromatin and/or possibly serve as protein-binding motifs and interact with some centromere-specific-proteins. The recent study on human centromeres has shown that the centromeric alpha satellite has an intrinsic tendency to form secondary structures including hairpins, but also that the binding of the CENP-B protein to the satellite repeats promotes formation of sub-micron-sized DNA loops, which are important for centromere stability and the maintenance of its position (Chardon et al. 2022). Conservation of inversion points in *T. freemani* centromeric satDNA arrays suggests that TfSat01 and TfSat03 macro-dyad symmetries might have some similar functional role.

The pronounced dyad symmetries also characterize the centromeric satDNAs of other *Tribolium* species, which have more complex monomeric units. In *Tribolium brevicornis*, the 1061 bp long monomer of the major satellite is composed of the two ∼470 bp long inversely oriented subunits (Mravinac et al. 2005). In *Tribolium madens*, a complex 704 bp long monomeric unit of the MAD2 satellite is also based on dyad symmetry (Ugarković, Durajlija, et al. 1996). In *Tribolium audax*, a related but much more complex monomer of the TAUD2 satellite consists of two ∼700 bp long inverted subrepeats (Mravinac and Plohl 2010). All these *Tribolium* major satellites are present in wide centromeric regions of all chromosomes. Therefore, we conclude that dyad symmetry may indeed be a structural imperative and that secondary structures in *Tribolium* centromeres may be more important than the primary structure. The superior importance of secondary structures could thus explain the lack of similarity in nucleotide sequences between the major centromeric satDNAs of congeneric species. Recent work on two closely related mouse species proposed that rapid evolution of (peri)centromeric DNAs does not obstruct chromatin packaging and chromosome segregation if the satellites adopt DNA shapes recognized by conserved architectural proteins (Dudka et al. 2025).

In the study of satDNA and chromatin organization in mouse centromeres, a low percentage of direction changes in pericentromeric and centromeric satDNAs’ arrays were detected, and the authors commented that they might result from the inversion events (Packiaraj and Thakur 2024). In the *T. freemani* centromeres, however, the direction switches are too pervasive to be explained by inversion events alone. We believe that the frequent macro-dyad symmetries in the *T. freemani* centromeres not only represent a possible structural preference, but may also indicate the mechanisms that promote the proliferation of centromeric satellites. One of these mechanisms could be via eccDNA molecules. SatDNA-derived eccDNAs have been found in a variety of eukaryotic species (Cohen et al. 2003; Cohen et al. 2006; Navrátilová et al. 2008; Huang et al. 2021; Liu et al. 2023), including *T. castaneum* (Volarić et al. 2024). In *T. freemani*, we confirmed the presence of centromeric major and minor satDNAs in the eccDNA fraction, so we suggest that the eccDNAs propagate adjacent arrays of the two satellites, promoting the dyad symmetries present therein. Our conclusion is additionally supported by preserved direction changes in the major satDNA arrays and in the minor satDNA-based intercalary segments, highly conserved in sequence even among non-homologous chromosomes. In addition to eccDNA-mediated spread, 3D spatial interactions in the nucleus among distant satellite arrays have been proposed as a mechanism that enables spread of satDNAs between remote loci (Sproul et al. 2020). In *T. freemani*, we detected satDNA-rich (peri)centromeric heterochromatin clustering in the meiotic prophase, which was also observed in other *Tribolium* species (Žinić et al. 2000; Mravinac and Plohl 2010). Non-homologous centromere pairing during early meiotic prophase I has been reported in multiple organisms (Kurdzo and Dawson 2015), including associations of satDNA-rich centromeric regions of the X chromosome and different autosomes in mouse (Spangenberg et al. 2021). In the study of murine telocentric chromosomes, which revealed high sequence identity between non-homologous chromosomes, it was concluded that this reflects a mechanism of frequent recombinational exchange between non-homologous chromosomes possibly facilitated by close associations during meiotic prophase (Kalitsis et al. 2006). The clustering of the non-homologous centromeres of *T. freemani* in early meiosis and the conservation of direction switches between satDNAs arrays at non-homologous chromosomes are in line with this interpretation.

### Potential functional implications of dynamic satDNA evolution

One of the important findings of this work is that *T. freemani* and *T. castaneum* despite completely different major satDNAs share an orthologous organization in their centromeres. In addition to dyad symmetries, in both species the arrays of species-specific major satDNAs intermingle with arrays of orthologous minor satDNAs. The fact that the siblings’ centromeres are remarkably different in the major satDNAs but similar in the less copious satellites contradicts the intuitive assumption that the most abundant satDNAs are the key sequences of the functional centromeres. At this point, we are inclined to hypothesize that the orthologous minor satDNAs (TfSat03 and TfSat04 from *T. freemani*, and Cast7 from *T. castaneum*) could be centromere-competent sequences. An analogy can be drawn with the centromeres of *Mus musculus*, in which the more abundant major satellite forms the pericentromeric heterochromatin, while the less abundant minor satellite is centromeric (Joseph et al. 1989). Interestingly, the mouse Y chromosome centromere is based on a highly diverged minor satellite-like sequence (Pertile et al. 2009). Similarly, the *T. freemani* sex chromosome y_p_ does not harbor the major satDNA, but comprises y_p_-specific satDNA TfSat04, from the superfamily of minor satDNAs. It could be that the y_p_-specific variant of minor satellite is developed and maintained by intrachromosomal homogenization due to suppressed exchange interaction with other chromosomes, also suggested by the absence of TfSat01/TfSat03 arrays on the y_p_ chromosome.

Functional centromeres are usually determined epigenetically by the presence of centromere-specific variants of histone H3, CENH3 (Mellone and Fachinetti 2021). By analyzing the centromeres of *T. castaneum*, we found that the regions of major satDNA TCAST encompass the CENH3 metapolycentric domains, but also extend beyond the CENH3 distribution (Gržan et al. 2020). Our preliminary testing of *T. castaneum* CENH3 antibodies in *T. freemani* indicates that *T. freemani* has its own species-specific CENH3, consistent with the centromere paradox hypothesis, which justifies the rapid evolution of centromeric DNA and protein components even among closely related species (Henikoff et al. 2001). Detailed studies on centromere-specific proteins, such as CENH3 histones, should help to decipher which of the *T. freemani* satDNAs, major or minor, are actually involved in centromere function.

*T. freemani* and *T. castaneum* separated 14 Mya ago and represent the youngest species within the >100 Mya old genus (Hinton 1948; Ramesh et al. 2021). Although it is not known whether the two siblings coexist in the same habitats, they interbreed under laboratory conditions and produce infertile F_1_ hybrids (Nakakita et al. 1981), indicating postmating reproductive isolation (Wade and Johnson 1994). Considering that the satDNAs are by far the most abundant sequences in their genomes and thus represent the largest genomic difference between the two species, it is tempting to speculate whether they might play a role in reproductive isolation. In two congeneric catfish species, genome-wide satDNA divergence between the parental species, rather than chromosome number discrepancy, was indicated as contributing to the sterility of the hybrids (Lisachov et al. 2024). In hybrids of closely related *Drosophila* species, differences in satDNA composition between parental species have been shown to impede the proper clustering of pericentromeric satDNAs into chromocenters in interphase nuclei, thereby promoting hybrid incompatibility (Jagannathan and Yamashita 2021). To determine whether satDNAs discrepancies underlie the reproductive isolation between *T. freemani* and *T. castaneum*, functional studies similar to those in *Drosophila* are required, and the hybrids will play a key role in answering this question.

In conclusion, the comprehensive study of the *T. freemani* satellitome and the comparison with *T. castaneum* revealed a dynamic evolution of satDNAs that yielded the greatest differences between the siblings’ genomes. Dissimilar levels of conservation, organization, chromosomal locations and abundances point to different evolutionary dynamics and mechanisms of propagation to which individual satellites within a genome are subject. As for the centromeric satDNAs, we deciphered the origin of the completely different major satellites and uncovered the orthologous organization of the centromeric regions, which show macro-dyad symmetries and related minor satDNAs as a commonality between the siblings. These findings provide a foundation for future work that will address the role of *Tribolium* most prominent satDNAs in the context of functionality and speciation, in which *T. freemani-T. castaneum* hybrids will be of great assist. In addition, this work serves as a good reference point for satellitome analyses of other *Tribolium* species, which should further improve the understanding of satDNA evolution in satDNA-rich genomes. We hold that satellite-abundant non-model organisms such as *Tribolium* can be very useful in tackling questions about functional and evolutionary implications of satDNA behavior, and our further research will go in this direction.

## MATERIAL & METHODS

### Insect material

The *T. freemani* beetles, originally obtained as a starter culture from the USDA Agricultural Research Service (Manhattan, KS, USA) in 2015, were propagated in the laboratory in whole wheat flour in a darkened incubator at 27⁰C and 50-70% relative humidity. The beetles were subcultured every 4-6 weeks.

### Genome assemblies

For the annotations of the repeats and the analyzes of the organization of the satDNA arrays, we used the high-quality chromosome-level genome assemblies that we recently generated: Tfree1.0 (NCBI GenBank accession number GCA_939628115.1) and TcasONT (ENA accession number GCA_950066185.1). The *T. freemani* genome assembly Tfree1.0 (262.9 Mb) is based on the PacBio HiFi reads (Volarić et al. 2022), and is currently designated as the *T. freemani* reference genome in the NCBI genome database. The *T. castaneum* genome assembly TcasONT (191 Mb) is based on Nanopore long-read sequencing and is the latest *T. castaneum* assembly, that has been significantly improved in repetitive regions (Volarić et al. 2024).

### DNA isolation and whole-genome sequencing

Total genomic DNA was extracted from 50 mg of snap-frozen adult *T. freemani* beetles (10 pooled male and female individuals) using the DNeasy Blood and Tissue Kit (Qiagen, Hilden, Germany). The isolated genomic DNA was quantified with Qubit 2.0 DNA HS Assay (ThermoFisher, Massachusetts, USA) and quality was assessed by Tapestation genomic DNA Assay (Agilent Technologies, California, USA). DNA was sent to the sequencing service provider Admera Health (South Plainfield, USA), where library preparation was done using KAPA Hyper Prep Kit (Roche, Switzerland). Illumina® 8-nt unique dual-indices were used to mitigate index hopping. Whole-genome sequencing (WGS) performed on an Illumina NovaSeq X Plus 10B platform (Illumina, California, USA) yielded 2×35,516,583 paired-end reads (2×151 nt). The resulting 10.7 Gb of sequenced data corresponds to approximately 35-fold coverage of the *T. freemani* genome. We have deposited the raw sequencing data in the NCBI Sequence Read Archive (SRA) database under the BioProject study accession number PRJNA1179347.

### SatDNA mining using graph-based clustering

The identification of *T. freemani* satDNAs from the unassembled short Illumina reads was performed by graph-based sequence clustering with the TAREAN pipeline (Novák et al. 2017). The raw reads were quality-checked and preprocessed using the RepeatExplorer2 tools on the Galaxy web server (https://repeatexplorer-elixir.cerit-sc.cz/galaxy/). The interlaced reads were randomly subsampled to reduce the size of the input dataset in order to achieve a low genome coverage required for TAREAN analyses. We used six randomly subsampled sets with 175,000 to 1,500,000 input reads, corresponding to genome coverages of 0.02–0.7x (**Suppl. Table S1**). Since the major satDNA TfSat01 accounts for 31% of the *T. freemani* genome, its high proportion considerably limits the number of reads that can be processed with the TAREAN tool, as determined in the first test analysis (**Suppl. Table S1,** T1 analysis). For this reason, in the subsequent analyses we used the input sets from which we excluded the TfSat01-containing reads (**Suppl. Table S1,** analyses T2-T6). A custom database was created containing the consensus sequences of the high and low putative satDNAs from all six TAREAN analyses, and blast search of the custom database was performed to exclude duplicates. The consensus sequences of the TAREAN satDNA candidates were mapped on the *T. freemani* genome assembly Tfree1.0, as described in the “Repeat annotations” paragraph below. If a candidate was found tandemly repeated in an array of at least five consecutive monomers, it was declared as satDNA. The consensus sequences of the 135 declared satDNAs are deposited in the NCBI GenBank under the accession numbers PQ553299-PQ553433. The satDNA consensus sequences were BLAST-searched against NCBI GenBank database (Clark et al. 2016) using megablast, discontiguous megablast, and blastn algorithms. The satDNA consensuses were screened against GIRI Repbase, the database of repetitive DNA elements (Bao et al. 2015) using the CENSOR tool.

### SatDNA bioinformatic analyses

The Geneious Prime 2023.2.1 package (Biomatters Ltd, New Zealand) was used for the basic analyses of satellite repeats such as monomer length, A+T composition, mutual pairwise similarities, and potential secondary structures. All other analyses were performed as described below, and the R scripts used in the analyses are available as Supplementary Code at Figshare (figshare.com/s/d9b5c22dcbe842f27a8d).

#### Repeat annotations

The annotation of the *T. freemani* satDNA consensus sequences in the *T. freemani* genome assembly Tfree1.0 and the *T. castaneum* genome assembly TcasONT was performed using NCBI’s stand-alone BLAST algorithm and the R programming language (R Core Team) with the metablastr package (Benoit and Drost 2021). The criteria for filtering the detected satDNAs in the genome were set to 70% percent identity and 70% query sequence coverage. The detected satDNA loci were converted to GFF files to facilitate visualization and extraction. The annotations of satDNA repeats are available as Supplementary Data S6.

#### Identification of satDNA arrays

SatDNA arrays were detected by sorting all hits for a given satDNA in the BLAST results table and checking whether another monomer of the same satDNA was found within a genomic distance of one monomer length. If a monomer was detected, the array was extended and the step was repeated until the array could no longer be extended (i.e. there were no repeats within a monomer length). This was particularly applied when searching for satDNAs arrays with ≥5 consecutive repeats or scattered organization. The annotations of satDNA arrays are available as Supplementary Data S6.

#### Genomic environment

Following array creation, the flanking regions of each array were examined for the presence of genes, transposable elements and satDNAs discovered in the previous step. The used annotations for genes and transposable elements were adopted from the official *T. freemani* annotation set available in FigShare at 10.6084/m9.figshare.19682400. The annotations for satDNAs are available as Supplementary Data S6. The search space for hits was 10 kb +/- from the respective array start/end, and the statistics of resulting hits were calculated.

#### Long-range organization of centromeric satDNAs

The first step in analyzing the major satDNA TfSat01 was to revise its consensus sequence. Our TfSat01 consensus obtained by TAREAN analysis of WGS reads showed 9 nucleotide differences compared to GenBank entry X58539 (**Suppl. Fig. S5A**), which was based on five randomly cloned monomers (Juan et al. 1993). We evaluated the two consensuses and concluded that our TfSat01 consensus is more representative based on the following arguments. First, we BLAST-searched the two consensuses against the Tfree1.0 assembly (using >70% similarity criterium) and detected 4.6% more monomer copies with the TfSat01 consensus (228 060 for TfSat01, 218 083 for X58539). Second, the distribution of similarities between the detected monomers and a query consensus showed a shift to the higher values for the TfSat01 consensus (**Suppl. Fig. S5B**), suggesting that the TfSat01 sequence is more typical. Third, although TfSat01 is derived as a consensus, it is not an artificial sequence. In the Tfree1.0 genome assembly, we discovered 32 monomer copies that were 100% identical to the TfSat01 consensus, while no monomers had 100% identity to X58539. We deposited the TfSat01 sequence as an updated consensus in the NCBI GenBank database under the accession number PQ553299.

To visualize the centromeric organization of TfSat01, we used StainedGlass software (Vollger et al. 2022) with the default settings and manually extracted centromeric regions of *T. freemani* chromosomes comprising multi-megabase long TfSat01 arrays. To detect the inverted TfSat01 subarrays, the array detection step was repeated in a strand-dependent manner to create the “strand-specific” database. Subsequently, the “strand-specific” arrays were overlapped with total TfSat01 arrays (strand-independent) to determine the number of inversions per strand-independent array. To determine the difference between the lengths of “strand-specific” subarrays, the total length of the minus-strand subarrays was deducted from the total length of the plus-strand subarrays and the differences were plotted. After detection of TfSat01 arrays, the gaps among TfSat01 arrays were identified and examined for the presence of the TfSat03 satDNA using overlaps and existing TfSat03 annotations. Following the detection of TfSat03 monomers in the gaps of TfSat01 arrays, the central regions where the TfSat03 repeats change orientation were extracted and analyzed using graph networks as described below. The observed patterns of long-range organization of the two centromeric satellites, including the intermingled organization of the TfSat01 and TfSat03 arrays as well as the inversion points within the arrays for both satellites, were additionally verified and confirmed on the raw PacBio HiFi reads used to create the Tfree1.0 assembly (Volarić et al. 2022).

#### Sequence alignments and visualizations

All detected monomers of the 135 *T. freemani* satDNA were matched with the corresponding arrays to assign them unique IDs and then extracted. To find orthologs in the *T. castaneum* genome, the same processes of detection, array creation and extraction were repeated for *T. freemani* satDNAs in the TcasONT assembly. Large multiple sequence alignments were generated using MAFFT (Katoh and Standley 2013). The genomic distance matrices were calculated using the dist.dna function with the “F81” genomic distance model from the *ape* package (Paradis and Schliep 2019). Principal Component Analysis (PCA) was subsequently performed on the distance matrix using the FactoMineR package (Lê et al. 2008) and visualized with ggplot2 (static plots) and plotly (dynamic HTML plots). Network visualizations were created from the distance matrices using the networkD3 package (available at https://cran.r-project.org/web/packages/networkD3/ index.html), by finding the 5 closest neighbors of each sequence in the alignment.

### Revision of fLG7 chromosome sequence in the *T. freemani* genome assembly Tfree1.0

The reference genome of *T. freemani*, Tfree1.0 (GCA_939628115.1) comprises the sequences of nine autosomes (fLG2-fLG10) and sex chromosome X (fLGX), but the sex chromosome y_p_ is not assembled (Volarić et al. 2022). According to the Tfree1.0 assembly, the satDNA TfSat04 is located at the end of chromosome fLG7, but in this work we established by *in situ* hybridization that the satDNA TfSat04 is y_p_-specific (**Fig. 4C**). Given the inconsistency between *in situ* and *in silico* localization of TfSat04, we revisited the contigs that we used to build the Tfree1.0 assembly (Volarić et al. 2022). We found that the longest continuous TfSat04 array is part of the ptg000052l contig, located distally on chromosome fLG7 and connected to the rest of the chromosome by an (N)_100_ placeholder representing an assembly gap (**Suppl. Fig. S20**). The ptg000052l contig ends in TfSat04 repeats and has one of the lowest location confidence scores in the assembly (Supplementary Table S3 in (Volarić et al. 2022)), so we hold that a slight similarity to TfSat03 caused the contig to be misaligned to fLG7. After reviewing its sequence in detail, we conclude that the ptg000052l contig is a candidate for the non-assembled y_p_ chromosome and support this conclusion with several findings. First, in addition to the longest arrays of TfSat04, this contig also harbors the longest array of the low-copy-number satDNA TfSat07 (**Suppl. Fig. S20**), and by *in situ* hybridization we indeed detected the strongest TfSat07 FISH signal on the y_p_ chromosome (**Suppl. Fig. S3C**). Second, given that assemblies in repetitive regions can be inaccurate, we examined the 24 Gb of the raw PacBio HiFi reads used for the Tfree1.0 assembly to analyze gene dosage. Gene dosages for 32 genes annotated on the ptg000052l contig and 2000 randomly selected genes annotated on different autosomes were estimated by mapping the genes at the raw reads with minimap2 (Li 2018). Notably, 32 genes annotated on the ptg000052l contig were represented in the raw reads on average two times less frequently compared to 2000 randomly selected genes (**Suppl. Table S10**). Third, regarding the contig ends, one side of the ptg000052l terminates in ∼18 kb TfSat04 array, while the other side ends in a 1.2 kb (TCAGG)_243_ array (**Suppl. Fig. S20**), which is the *Tribolium* telomeric sequence (Mravinac et al. 2011). The repetitive endings suggest that the repeats at both ends extend further, but the ptg000052l contig with its size of 2.2 Mb probably accounts for the majority of the y_p_ sequence, the smallest chromosome of *T. freemani*.

### SatDNA probes

To localize satDNAs on the chromosomes of *T. freemani* by fluorescence *in situ* hybridization (FISH) or to prove their presence in the extrachromosomal circular DNA (eccDNA) fraction, specific DNA probes were prepared. Specific probes were generated by PCR amplification of fragments of a satDNA of interest from the genomic DNA, and subsequent cloning into a plasmid vector. A modified version of Primer3 2.3.7, implemented in Geneious Prime 2023.2.1 package (Biomatters Ltd, New Zeland), was used to design specific primers for each satDNA studied. The sequences of primer pairs and their optimal annealing temperatures are specified in **Suppl. Table S11**. Reaction mixtures contained 10 ng genomic DNA, 0.2 μM of each specific primer, 0.2 mM dNTP mix, 2.5 mM MgCl2, 0.25 U GoTaq G2 Flexi DNA polymerase, and 1x Colorless GoTaq Flexi Buffer (Promega, USA). PCR amplification consisted of predenaturation at 94 ⁰C for 3 min, 35 amplification cycles (denaturation at 94 ⁰C for 10 s, annealing at optimal T_a_ for 10 s, extension at 72 ⁰C for 10 s), and a final extension at 72 ⁰C for 5 min. Cloning of satDNA fragments into the pGEM-T Easy plasmid vector (Promega, USA) and transformation of *Escherichia coli* XL10-Gold Ultracompetent Cells (Agilent Technologies, USA) were performed according to the manufacturer’s instructions. Putative positive clones were selected by blue-white color screening for recombinant plasmids, and insert lengths were checked via colony-PCR amplification using the vector-specific primers M13F and M13R-40. Finally, insert sequences of the clones used for satDNA probes were verified by Sanger sequencing (Macrogen Europe BV, the Netherlands). To cover the variability of monomers within a satellite DNA, a mixture of several clones was generally used to label each of the satDNA probes. SatDNA probes were labeled with biotin-16-(5-aminoallyl)-dUTP or aminoallyl-dUTP-Cy3 (Jena Bioscience, Germany) by PCR amplification using specific primers and cloned inserts as a template under conditions specified above. The ratio of labeled dUTP and dTTP in the labeling reaction was 1:2.

### Chromosome preparations and fluorescence *in situ* hybridization (FISH)

For FISH analyses, gonads isolated from *T. freemani* pupae were used to prepare chromosome spreads using the squash method. Freshly isolated testes or ovaries were incubated in 10 μg/ml colcemid (Roche, Switzerland) for 1 h. Hypotonic treatment was carried out in 0.075 M KCl for 5-15 min, followed by fixation in a solution of absolute ethanol:acetic acid (3:1) for 15-30 min. The gonads were dissected on a slide in a drop of 45% acetic acid, covered with a coverslip and squashed. Slides with coverslips were immersed in liquid nitrogen for 30 s and the coverslips immediately removed with a razor blade. Slides were air-dried and stored at -80 ⁰C until use. Prior to *in situ* hybridization, slides with chromosome spreads were subjected to the following treatments: incubation with RNase A (100 μg/mL) for 1 h at 37 ⁰C, incubation in 0.01% pepsin in 10 mM HCl for 10 min at 37 ⁰C, incubation in 2.7% formaldehyde solution in PBS for 10 min at room temperature, dehydration through an ice-cold ethanol series (70% → 90% → 100%, 3 min each), denaturation in 70% formamide at 70 ⁰C for 2 min, and final dehydration through an ice-cold ethanol series. Labeled satDNA probes (100-200 ng per slide) were first lyophilized (in double-color FISH experiments, probes for different satDNAs were mixed before lyophilization), then dissolved in 15 µl of hybridization buffer (60% formamide, 40% DeSO_4_ buffer) and denatured at 75 ⁰C for 5 min. After cooling on ice, denatured probes were applied to chromosome slides and incubated in a moist chamber at 37 ⁰C for overnight hybridization. Posthybridization washes were performed at 37 ⁰C in 50% formamide in 2xSSC at 37 ⁰C. While Cy3-labeled probes were ready for direct detection after posthybridization washes, biotin-labeled probes were visualized with fluorescein avidin D and biotinylated anti-avidin D system (Vector Laboratories, USA) using signal amplification. Signal amplification was achieved through three successive incubations in: 1) 1:500 dilution of fluorescein avidin D, 2) 1:100 dilution of biotinylated anti-avidin D, and 3) 1:2000 dilution of fluorescein avidin D. Finally, slides were counterstained in 4’,6-diamidino-2-phenylindole (DAPI) solution and embedded in Mowiol 4–88 mounting medium (Sigma-Aldrich, USA).

### Confocal microscopy and image analyses

Slides were examined with a Leica TCS SP8 X confocal laser scanning microscope, equipped with a HC PL APO CS2 63/1.40 oil objective, a 405 nm diode laser and a supercontinuum excitation laser (Leica Microsystems, Germany). Images were captured separately for each fluorochrome and processed with LASX Office 1.4.7 28921 (Leica Microsystems, Germany), ImageJ (Schneider et al. 2012) and Adobe Photoshop CS5 (Adobe Systems, USA), using only functions affecting the entire image equally. For each satDNA, at least 10 metaphase spreads from 3-10 independent experiments were analyzed.

### Extrachromosomal circular DNA isolation and two-dimensional agarose gel electrophoresis

To test the presence of satDNAs in the extrachromosomal circular DNA (eccDNA) molecules, total genomic DNA was isolated from 920 mg of mixed male and female larvae and adults of *T. freemani* according to the procedure described in (Volarić et al. 2021). DNA quantity was determined with a Qubit 4 fluorometer using Qubit dsDNA BR Assay Kit (Invitrogen, USA). In order to remove as much linear double-stranded DNA (dsDNA) as possible from the isolated DNA, the isolated DNA was treated mechanically and enzymatically. First, 20 μg of the isolated DNA was passed through a 0.33 mm syringe needle 30 times to shear the linear DNA. The sheared DNA was then treated with Exonuclease V, which degrades linear dsDNA in both 3’ to 5’ and 5’ to 3’ directions leaving intact circular DNA. 20 μg of sheared DNA was digested with 200 U Exonuclease V (New England Biolabs, USA) at 37 ⁰C overnight, and digestion was stopped by 11 mM EDTA pH 8.0 and incubation at 70 ⁰C for 30 min. Prior to two-dimensional (2D) electrophoretic separation, DNA was cleaned using the Monarch® PCR & DNA Cleanup Kit (New England Biolabs, USA). Electrophoresis in the first dimension was performed in 0.7% ethidium bromide (EtBr)-free agarose gel in 1xTBE buffer at low voltage (0.7 V/cm) for 18 h. The first-dimension gel was then stained in 1x TBE with 0.3 µg/ml EtBr for 1 h. The entire gel strip of interest from the 1D agarose gel was excised, placed at a 90° angle to the direction of electrophoresis on a casting tray and poured with 1.5% agarose containing 0.3 µg/ml EtBr. The second-dimension electrophoresis was performed in 1xTBE containing 0.3 µg/ml EtBr at a higher voltage (4 V/cm) for 3 h. Both electrophoreses were run at room temperature.

### Southern blot hybridization

After 2D electrophoresis, agarose gels with separated DNA were agitated in 0.25 M HCl for 30 min and then in 0.4 M NaOH for 30 min. Depurinated and denatured DNA fragments were blotted onto a positively charged nylon membrane (Roche, Switzerland) overnight by capillary alkaline transfer in 0.4 M NaOH. Hybridization probes for TfSat01 and TfSat03 were prepared by PCR labeling using biotin-16-(5-aminoallyl)-dUTP (Jena Bioscience, Germany) and clones carrying specific satDNA fragments as described previously. Hybridization was performed in hybridization buffer (250 mM Na_2_HPO_4_ pH 7.2, 1 mM EDTA, 20% SDS, 0.5% blocking reagent) with agitation overnight at 68 ⁰C, while posthybridization washing was done in washing buffer (20 mM Na_2_HPO_4_, 1 mM EDTA, 1% SDS) at 65 ⁰C. Chemiluminescent detection was carried out using the streptavidin-alkaline phosphatase (streptavidin-AP) conjugate and CDP-Star substrate (Roche, Switzerland). The signals were visualized using the Alliance Q9 Mini imaging system (Uvitec, UK). To hybridize the same blot with another probe, the first probe was stripped by washing the blot in stripping buffer (0.2 M NaOH, 0.1% SDS) at 42 ⁰C for 2 x 15 min and then rinsed 3 x 5 min at room temperature in 2xSSC. After stripping and between hybridization experiments, blots were stored in 2xSSC at 4 ⁰C to keep them wet. The stripping efficiency was tested by re-application of the streptavidin-AP conjugate and detection with the CDP-Star. To prove that stripping of the first probe does not affect hybridization with the second probe, multiple experiments were performed in which the order of the hybridization probes was changed.

## DATA AVAILABILITY

The raw Illumina sequencing data of the *T. freemani* genome are deposited in the NCBI Sequence Read Archive (SRA) database under the BioProject study accession number PRJNA1179347. The consensus sequences of the 135 *T. freemani* satDNAs are deposited in the NCBI GenBank under the accession numbers PQ553299-PQ553433. All scripts used for the analyses in this study are available as Supplementary Code at Figshare (figshare.com/s/d9b5c22dcbe842f27a8d). All supplementary figures, tables and data are available as Supplementary Material at Figshare (figshare.com/s/d9b5c22dcbe842f27a8d).

## AUTHORS’ CONTRIBUTIONS

Conceptualization - BM; Investigation - DV, EDS, MV, LH; Formal analysis - DV, EDS, MV, TVZ, BM; Visualization - DV, EDS, MV, BM; Resources - NM, BM; Funding acquisition and project administration - BM; Supervision - BM; Writing - original draft: BM; Writing review and editing - DV, EDS, MV, NM, BM. All authors read and approved the final manuscript.

## SUPPLEMENTARY MATERIAL

**Supplementary Figure S1.** Consensus sequence alignments of satDNAs belonging to four superfamilies (SF1-SF4). The attached matrices show the percentage of pairwise identity between the compared sequences. For the superfamily SF1 (**A**), the comparison between the satellite TfSat04 and the three subunits belonging to the satDNA TfSat03 is presented.

**Supplementary Figure S2.** Alignments of the subunits of satDNA TfSat03 (**A**), TfSat39 (**B**), TfSat41 (**C**), TfSat76 (**D**), and TfSat77 (**E**), whose monomers are based on higher-order-repeats (HORs). Schematic representations show the HOR structure, the lengths of the subunits and their pairwise similarities.

**Supplementary Figure S3.** Localization of TfSat05 (**A**), TfSat06 (**B**), TfSat07 (**C**), TfSat08 (**D**), TfSat09 (**E**) and TfSat10 (**F**) satDNAs on *T. freemani* chromosomes (2n=20), determined by fluorescence *in situ* hybridization. Chromosomes were stained with 4′,6-diamidino-2-phenylindole (DAPI) and hybridized with satDNA probes visualized by fluorescein isothiocyanate (FITC). For each satDNA localization, chromosomes are shown in black-and-white (first panel) and in DAPI-stained version (second panel). The position of a satDNA-specific probe is indicated by green fluorescence in the third panel, while the fourth panel shows a merge of DAPI-stained chromosomes and FITC-vizualized satDNA probes. For satDNA TfSat07 (**C**), the position of the signals on the male chromosome y_p_ is indicated by an arrow. The scale bar corresponds to 3 µm.

**Supplementary Figure S4.** Distances of genes, satellite DNAs (satDNAs) and transposable elements (TEs) to *T. freemani* satDNAs arrays (containing ≥5 consecutive monomers) in their 10-kb flanking regions. Each dot represents the mean distance of annotated sequences of interest for each satDNA. The x-axis indicates the mean position value in the 10-kb flanking region, with negative and positive values indicating the left and right flanking regions, respectively. The mean distance values were calculated based on the data provided for individual arrays in Supplementary Table S5.

**Supplementary Figure S5.** Comparison of the consensus sequences TfSat01 (defined in this work) and GenBank entry X58539 (Juan *et al*., 1993). **A)** The alignment of the consensus sequences. The nucleotide differences between the compared consensuses are colored, and dashes indicate alignment gaps. **B)** Distribution of nucleotide sequence similarities between monomers annotated in the *T. freemani* genome assembly Tfree1.0 and the consensus sequence TfSat01 (left panel) and the consensus X58539 (right panel). The BLAST search was done by using the criteria of >70% sequence similarity and >70% query coverage. The black arrow points to the monomer group that has 82% similarity to the TfSat01 consensus.

**Supplementary Figure S6.** TfSat01 multi-megabase regions in the Tfree1.0 assembly visualized with StainedGlass (Vollger *et al*., 2022). The StainedGlass analysis was performed using 10 kb windows, and the sequence identity values are shown in the color histogram along each sequence identity heatmap. Two discovered multi-megabase regions (region 1 and region 2) are shown for chromosome fLG9. The lengths and positions of the multi-megabase regions are listed in Supplementary Table S8.

**Supplementary Figure S7.** Density plots of the differences between the length of the direct and inverted subarray within the 1793 TfSat01 arrays distributed at seven *T. freemani* chromosomes (fLG2-fLG9). The X-axis shows the difference in the length of directly and inversely oriented subarrays within an array. The relative abundance of subarray length differences in the graph is indicated by color gradient.

**Supplementary Figure S8.** Organization of the six most frequent inversion segments (26 bp, 106 bp, 132 bp, 149 bp, 187 bp, and 200 bp long) in which TfSat01 monomers change their orientation within the TfSat01 arrays. **A)** A schematic showing the position of the inversely oriented truncated TfSat01 monomers in the inversion segments. The labels “DIR” and “INV” indicate the direct and inverted orientation of the truncated monomers within the inversion segment. **B)** Alignment of the truncated TfSat01 monomers in the inversion segments. The sequence “TfSat01_direct_vs_inverted” represents directly and inversely oriented TfSat01 consensus monomers, according to which the truncated monomers of the inversion site are aligned. **C)** Potential secondary structures of six inversion segments predicted by the RNAfold tool (Lorenz *et al*., 2011) using DNA Matthews 1999 energy model. The minimum free energies of the structures are given in parentheses. **D)** List of chromosomes on which a particular inversion segment is present. The alignments of sequences belonging to the particular inversion segments are shown in Supplementary Data S1.

**Supplementary Figure S9.** The sequence of the central region in the intercalary segment where the degenerate TfSat03 copies change orientation. **A)** Schematic representation of the central region with the inverted segments marked by red and green arrows. **B)** Potential secondary structure of the central region predicted by the RNAfold tool (Lorenz *et al*., 2011) using DNA Matthews 1999 energy model (the minimum free energy of the structures = -6.06 kcal/mol).

**Supplementary Figure S10.** Graph network of ∼110 bp long central regions from 1577 intercalary segments in which TfSat03 monomers change their orientation. The graph illustrates the sequence similarity relationships based on the alignment presented in Supplementary Data S2. Each dot represents one sequence, and the color of the dot indicates the chromosome on which the sequence is located. The color legend indicates individual chromosomes. The cluster, which consists of 100% identical sequences originating from different chromosomes, is marked with a black square and shown enlarged in Figure 3D. An interactive representation of the graph network is given in the Supplementary Data S3.

**Supplementary Figure S11.** The alignment between TfSat01, TfSat02 and TCsat15 consensus sequences. Identities between the compared sequences are indicated by dots (.), and dashes (-) indicate alignment gaps. The box indicates the 166 bp segment within the TfSat02 and TCsat15 consensuses that corresponds to the TfSat01 sequence. The blue arrow shows the position and orientation of the TfSat01 sequence.

**Supplementary Figure S12.** TCAST transposon-like element structure and genome distribution. **A)** Schematic representation of the TCAST transposon-like element with the indicated positions of a 367 bp long segment corresponding to the TCAST satDNA monomer (TCAST) and its 143 bp long truncated version (TCAST***). The inverted orientation of the segments is indicated by the direction of the red arrows. The terminal inverted repeats (TIR) of the DNA transposon-like element are indicated by gray arrows. **B)** Distribution of the 57 copies of the TCAST transposon-like element annotated in the *T. castaneum* assembly TcasONT (upper panel) and 84 copies annotated in the *T. freemani* genome assembly Tfree1.0 (lower panel). **C)** Maximum likelihood tree of the individual copies of the TCAST transposon-like element detected in the *T. castaneum* assembly (red) and in the *T. freemani* assembly (blue). The tree was inferred in iTOL with nodal supports based on 100 bootstrap replicates (the values ≥80 are shown).

**Supplementary Figure S13.** Relationships between *T. freemani* satDNAs TfSat03 and TfSat04, and *T. castaneum* satDNA Cast7. In the consensus sequences’ alignment of Cast7 and TfSat04 monomers and TfSat03 subunits A, B and C, differences between the compared consensuses are colored and dashes indicate alignment gaps. The attached matrix shows the percentage of pairwise identity between the compared sequences.

**Supplementary Figure S14.** Alignments of the satDNAs *T. freemani* satDNAs (TfSat) and their orthologs among the known *T. castaneum* satDNAs (Cast and TCsat). Pairwise identities between the consensus sequences are indicated in the parentheses.

**Supplementary Figure S15.** Relationships between the *T. freemani* satDNAs and their *T. castaneum* orthologs associated with DNA transposons from the Rehavkus superfamily. **A)** Phylogenetic relationships between the TfSat15 and TCsat23 repeats present in the inverted termini of the Rehavkus-1_TC DNA transposon. The schematic of the Rehavkus-1_TC is shown in the center, while the relationships between the *T. freeemani*/*T. castaneum* satDNA repeats from the inverted termini and the relationships between the central part of the transposon sequence are shown in the upper and lower panels, respectively. The sequences of *T. freemani* are represented by blue dots, while red dots indicate *T. castaneum* sequences. The maximum likelihood trees were inferred in iTOL with nodal supports based on 100 bootstrap replicates. **B)** Phylogenetic relationships between the TfSat23 and its orthologous *T. castaneum* repeats present in the inverted termini of the Rehavkus-3_TC DNA transposon. The same description of the tree applies as for the previous one. **C)** The principal component analysis (PCA) of the TfSat25 repeats present in the Rehavkus-like element in *T. freemani* and the orthologous repeats of the *T. castaneum* satDNA TCsat12 present in the 80.5 kb long array on *T. castaneum* chromosome LG4. In the PCA plot, the *T. freemani* repeats are indicated by blue dots, while the *T. castaneum* repeats are indicated by red dots. Schematic on the right illustrates the different organization of the orthologous repeats in the two sibling species.

**Supplementary Figure S16.** Comparison of the 10-kb flanking regions of the orthologous satDNAs TfSat08 and TCsat17 reveals the syntenic locus where the satDNAs are located in *T. freemani* (on chromosome fLG3) and *T. castaneum* (on chromosome LG3). The interspecific sequence similarities between the flanking regions are indicated within up-down arrows.

**Supplementary Figure S17.** Organization of the *T. freemani* satDNA TfSat06. **A)** The principal component analysis of the TfSat06 repeats from different *T. freemani* chromosomes. Dots represent repeats colored according the chromosome (fLG) of origin. **B)** Schematic representation of the organization of the Rehavkus-like element whose inverted termini harbor the TfSat06 repeats (represented by green arrows). The table reports the segmental similarities of the central part of the Rehavkus-like element with the repetitive elements deposited in the Repbase database. The similarities refer to the segments (purple blocks in the diagram) whose positions are listed in the table together with the similarities and alignment scores.

**Supplementary Figure S18.** Principal component analysis (PCA) of the orthologous satDNAs TfSat22 and Cast9. The upper panel shows intense mixing of *T. freemani* TfSat22 repeats (blue dots) and *T. castaneum* Cast9 repeats (red dots). Interchromosomal homogenization within the species is shown in the lower panels (*T. freemani* left, *T. castaneum* right), with the repeats of individual chromosomes colored according to the attached color legend.

**Supplementary Figure S19.** Organization of the *T. freemani* low-copy-number satDNAs TfSat54 (**A**) and TfSat97 (**B**), showing intraarray homogenization. The positions of the arrays on the chromosomes are indicated on the schematics (the size of the chromosomes is indicated on the left). Clusters formed by repeats of the same array are circled in the PCA plots. Red dots represent orthologous copies detected in the sibling species *T. castaneum*.

**Supplementary Figure S20.** The position of the ptg000052l contig (green rectangle in the center panel) on the *T. freemani* chromosome fLG7 in the Tfree1.0 assembly. The top panel shows an enlarged view of the junction between the pg000052L contig and the rest of the fLG7 chromosome sequence through the (N)_100_ placeholder. The lower panel shows the ptg000052l contig with the indicated positions of satDNA TfSat04 (magenta arrows), satDNA TfSat07 (brown arrows) and telomeric repeats TCAGG (dark green arrows).

**Supplementary Table S1.** The output of six clustering analyses (T1-T6) performed by the TAREAN pipeline using Illumina paired-end whole genome sequence reads of the flour beetle *T. freemani*. With the exception of analysis T1, all other analyzes (T2-T6) used sets of input reads from which reads corresponding to the major satDNA (Juan *et al*., 1993) were excluded. The genome coverage was calculated based on the estimated T. freemani genome size of 320 Mb (Volarić *et al*., 2022).

**Supplementary Table S2.** Consensus sequences and GenBank accession numbers of the 135 *T. freemani* satDNAs identified in this work.

**Supplementary Table S3.** Main characteristics of the 135 *T. freemani* satDNAs identified in this work. The last two columns specify orthologous satDNAs from the sibling species *T. castaneum* found by BLAST searches against the NCBI GenBank database and repetitive elements from the Repbase database with which *T. freemani* satDNAs showed partial similarities when searched with the CENSOR tool.

**Supplementary Table S4.** Number of satDNA monomers annotated in the Tfree1.0 genome assembly using >70% sequence similarity criterion.

**Supplementary Table S5.** Distance of nearest annotated genes, satellite DNAs (satDNA) and transposable elements (TE) in the flanking 10-kb regions (left and right) for individual arrays of the *T. freemani* satDNAs. The arrays are defined as stretches containing ≥5 consecutive monomers. The array ID name indicates the fLG chromosome (beginning of the name) and the position on the chromosome (end of the name) on which the array is located. Positions and distances are given according to the annotations in the Tfree1.0 genome assembly (Volarić *et al*., 2022).

**Supplementary Table S6.** Summary data on the presence of genes, satDNAs and transposable elements (TE) in the 10-kb flanking regions of the *T. freemani* satDNAs. The summary data are based on the data given for individual arrays in Supplementary Table S5. “Yes” indicates that at least one array has an annotated sequence of interest in its 10-kb surrounding region (left or right), whereas “No” indicates that no array has an annotated sequence of interest in the 10-kb surrounding regions.

**Supplementary Table S7.** Average distance (mean and median) of genes, satellite DNAs (satDNAs) and transposable elements (TEs) from *T. freemani* satDNA arrays (with ≥5 consecutive monomers) analyzed in their 10-kb flanking regions (left and right). Average distances were calculated based on the data given for individual arrays in Supplementary Table S5.

**Supplementary Table S8.** Lengths and positions of the TfSat01 multi-megabase regions on individual chromosomes in the Tfree1.0 assembly.

**Supplementary Table S9.** The number of the subarrays within a continuous TfSat01 array. The second column indicates the number of TfSat01 arrays annotated in the Tfree1.0 assembly, which contain the number of subarrays specified in the first column.

**Supplementary Table S10.** Quantitates of 32 genes from the ptg000052L contig and 2000 randomly selected genes from the Tfree1.0 assembly in 24 GB of raw data obtained by PacBio HiFi sequencing of the *T. freemani* genome. The occurrence of the genes was determined by mapping their sequences against HiFi reads using the minimap2 tool.

**Supplementary Table S11.** The sequences of *T. freemani* specific satDNA primers used in this study. The optimal annealing temperature (T_a_) for each primer pair is indicated.

**Supplementary Data S1.** Alignments of segments in which TfSat01 monomers change their orientation within the TfSat01 arrays. Shown are the alignments for the six most common segments (26 bp, 106 bp, 132 bp, 149 bp, 187 bp and 200 bp long). The names of the aligned sequences indicate the *T. freemani* fLG chromosome (beginning of the name) and the position on the chromosome (end of the name) on which the sequence is located.

**Supplementary Data S2.** The alignment of 1577 sequences of the ∼110 bp long central region of intercalary segments in which TfSat03 monomers change their orientation. The names of the aligned sequences indicate the *T. freemani* fLG chromosome (beginning of the name) and the position on the chromosome (end of the name) on which the sequence is located.

**Supplementary Data S3.** Interactive graph network of ∼110 bp long central regions from 1577 intercalary segments in which TfSat03 monomers change their orientation. The graph illustrates the sequence similarity relationships based on the alignment presented in Supplementary Data S2. Each dot represents one sequence, and the color of the dot indicates the *T. freemani* fLG chromosome on which the sequence is located. The color legend indicates individual chromosomes. The position of the sequence on the chromosome is revealed by touching a dot with the mouse pointer. The relationships between the individual sequences within the network can be examined by zooming in/out.

**Supplementary Data S4.** Alignments of *T. freemani* satDNAs (TfSat) monomers annotated in the *T. freemani* Tfree1.0 genome assembly and their orthologous copies annotated in the *T. castaneum* TcasONT assembly. For each annotated monomeric repeat, the beginning of the name (Tfree/Tcas) indicates the species in whose assembly the repeat is annotated. The designations fLG and LG in the names of the repeats indicate the chromosome and the position at which the repeat is annotated (fLG for *T. freemani* chromosomes, LG for *T. castaneum* chromosomes).

**Supplementary Data S5.** Principal component analyses (PCAs) of aligned *T. freemani* satDNAs (TfSat) repeats annotated in the *T. freemani* Tfree1.0 genome assembly and their orthologous copies annotated in the *T. castaneum* TcasONT assembly. The PCA plots are based on the alignments presented in Supplementary Data S4. The dots represent monomer repeats colored according to the species of origin (*T. freemani* in blue, *T. castaneum* in red). By moving the mouse pointer over a dot, the chromosomal location of the annotated monomer is displayed indicating a chromosome (fLG for *T. freemani*, LG for *T. castaneum*) and repeat’s position on a chromosome.

**Supplementary Data S6.** Annotations of *T. freemani* stDNA monomers and arrays annotated in the *T. freemani* genome assembly Tfree1.0.

